# Proteomic Insights into Breast Cancer Response to Brain Cell-Secreted Factors

**DOI:** 10.1101/2023.10.22.563488

**Authors:** Shreya Ahuja, Iulia M. Lazar

**Author notes:** **Correspondence:** Iulia M. Lazar, **E-mail:**.

## Abstract

The most devastating feature of cancer cells is their ability to metastasize to distant sites in the body. HER2+ and TN breast cancers frequently metastasize to the brain and stay potentially dormant for years until favorable conditions support their proliferation. The sheltered and delicate nature of the brain prevents, however, early disease detection and effective delivery of therapeutic drugs. Moreover, the challenges associated with the acquisition of brain biopsies add compounding difficulties to exploring the mechanistic aspects of tumor development, leading to slow progress in understanding the drivers of disease progression and response to therapy. To provide insights into the determinants of cancer cell behavior at the brain metastatic site, this study was aimed at exploring the early response of HER2+ breast cancer cells (SKBR3) to factors present in the brain perivascular niche. The neural microenvironment was simulated by using the secretome of a set of brain cells that come first in contact with the cancer cells upon crossing the blood brain barrier, i.e., endothelial cells (HBEC5i), astrocytes (NHA), and microglia (HMC3). Cytokine microarrays were used to investigate the secretome mediators of intercellular communication, and proteomic technologies for assessing the changes in the behavior of cancer cells upon exposure to the brain cell-secreted factors. The cytokines, growth factors and enzymes detected in the brain secretomes were supportive of inflammatory conditions, indicating a collective functional contribution to cell activation, defense, inflammatory responses, chemotaxis, adhesion, angiogenesis, and ECM organization. The SKBR3 cells, on the other hand, secreted numerous cancer-promoting growth factors that were either absent or present in lower abundance in the brain cell cultures, indicating that upon exposure the SKBR3 cells may have been deprived of favorable conditions for optimal growth. Altogether, the results suggest that the subjection of SKBR3 cells to the brain cell-secreted factors altered their growth potential and drove them toward a state of quiescence, with broader overall outcomes that affected cellular metabolism, adhesion and immune response processes. The findings of this study underscore the key role played by the neural niche in shaping the behavior of metastasized cancer cells, provide insights into the cellular cross-talk that may lead cancer cells into dormancy, and highlight novel opportunities for the development of metastatic breast cancer therapeutic strategies.

## Introduction

Brain metastasis accounts for more than half of the intracranial tumors in adults, making it one of the primary causes of brain cancer in patients [1]. Tumors of the lung, breast, and kidney, in addition to melanoma and colorectal cancer, have the highest propensity to spread to the brain, with approximately 80 % of cancers metastasizing to the cerebral hemispheres due to favorable blood flow patterns [1]. Moreover, recent statistics indicate that among the breast cancer patient cohort, the triple negative and HER2+ breast cancer subtypes are the most common to metastasize to the brain with a median incidence rate of 31 % and 32 %, respectively [2]. In order to colonize the brain, the breast cancer cells must breach the blood brain barrier (BBB), invade the neural matrix, and engage in a cross-communication of signaling molecules with the brain cells to evade the eliminatory signals from the host matrix. Crossing the BBB is a challenge in itself, as the endothelial cells of the BBB are linked together by tight junctions that restrict the paracellular movement of most large molecules and cells. Once the BBB has been breached, the tumor cells attach themselves to the brain microvasculature, a necessary process for extracting nourishment and proliferation cues [3]. In this stage, the cancer cells can stay dormant for years [4]. To create favorable growth conditions and further enable successful colonization, the cancer cells engage with the brain cells through direct physical interactions or via paracrine signaling. The neurons and astrocytes produce large amounts of extracellular matrix which accounts for ∼20 % of the CNS volume, alongside a broad spectrum of neural stimulants like trophic factors, neurotransmitters, cytokines, etc., that render a fertile ground for tumor outgrowth [5]. For example, the gap junction connexin Cx43 that interconnects the network of astrocytes has been found to also instigate cell survival and proliferative signaling in cancer cells [6]. Nevertheless, the brain also contains its own population of macrophages, the microglia, that survey the CNS for pathogenic insults and initiate immune responses. In this unfamiliar environment, the metastatic cancer cells face unique challenges, and, in response, express anti-inflammatory mediators that can suppress the immune activation of microglia and reprogram them to become tumor-supportive. Several studies have underscored the complex interactions that occur between tumor cells and their microenvironment, and that lead to the activation of processes and pathways that favor the development of metastases [7,8]. Very little is known, however, about the early molecular responses which are activated in cancer cells when they encounter the foreign neural niche. To gain insights into this process, the objective of this study was to explore the response of SKBR3/HER2+ breast cancer cells to factors expressed by three main cell types present in the neurovascular unit, i.e., microglia, astrocytes, and brain endothelial cells. The “neural niche” conditions were replicated by treating the cancer cells with conditioned media (CM) collected from *in vitro* cultured cells that contain secreted factors such as proteins, cytokines, growth factors, as well as extracellular vesicles (EVs) released by the cultured cells [9]. These components provide the ground for establishing the intercellular communication and promote processes that can accelerate the malignant growth of cells. In one such study, for example, the authors reported the presence of angiogenic factors FGF2 and VEGF in EVs shed by astrocytes in culture [10]. Astrocytes have been also found to interact with the invading tumor cells in early stages of brain metastases by secreting factors such as MMP9 and SDF1 that in turn support cancer progression [11]. Another group demonstrated that the astrocyte-conditioned medium contained exosomes with high levels of microRNAs which inhibit the expression of tumor suppressor PTEN in metastatic breast cancer cells, leading to aggressive tumor formation in the brain [12]. Together with astrocytes, microglia too secrete a variety of cytokines and factors that contribute to immunosuppression, increased invasiveness, angiogenesis, and cancer cell proliferation [13,14]. Furthermore, microglial EVs have been found to induce survival and provide metabolic support to other cells [15]. Owing to their role in promoting neurogenesis, maintenance of neuronal circuitry and blood brain barrier functions, microglia and astrocytes secrete neurotrophic factors to carry out neuroprotective and developmental functions in the brain [16–18]. Previous studies have highlighted the brain-derived neurotrophic factor (BDNF) released by astrocytes, which activates the tropomyosin-related kinase B (TrkB) receptor on metastasized breast cancer cells to promote cell proliferation and colonization of the brain [19]. Therefore, we hypothesized that the factors that are present in the CM collected from brain cells would alter the proliferative capacity of breast cancer cells and induce changes in the proteome that are suggestive of increased metastatic potential. To test our aforementioned hypothesis, we first explored the cytokine and growth factor profile of the conditioned medium collected from the three brain cell cultures by using a panel of capture antibodies spotted on a membrane-based cytokine array. Next, we used high-resolution mass spectrometry (MS)-based proteomics to measure the change in protein abundances between the CM-treated and non-treated SKBR3 cells. Finally, we mapped these changes to biological processes that are reflective of cancer growth and progression. This study provides preliminary, yet novel, insights into the early response of breast cancer cells exposed to *in vitro* conditions that mimic the neural niche. Our results suggest that the exposure of SKBR3 cells to neural niche factors may drive the cells into an initial state of quiescence, which is a possible result of inflammatory mediators and reduced levels of certain growth factors in the CM from brain cells.

## Experimental methods

### Supplies

Human HER2+ breast cancer cells (SKBR3), microglia (HMC3), brain endothelial cells (HBEC5i), and penicillin-streptomycin (PenStrep) solution were purchased from ATCC (Manassas, VA, USA). Normal human astrocytes (NHA), Clonetics AGM^TM^ Astrocyte Growth Medium BulletKit^TM^ (comprised of ABM-astrocyte basal medium and SingleQuots^TM^ growth factors, cytokines, and supplements pack), and astrocyte ReagentPack^TM^ subculture reagents were procured from Lonza (Morristown, NJ). Minimum essential medium (MEM), McCoy’s 5A (Modified) medium, DMEM/F-12, DMEM high glucose (HG), Dulbecco’s phosphate buffered saline (DPBS) and trypsin-EDTA were obtained from Gibco (Gaithersburg, MD). BenchMark Fetal Bovine Serum (FBS) was purchased from Gemini Bio Products (West Sacramento, CA). Endothelial cell growth supplements (ECGS) and all commonly used reagents (Na2VO3, NaF, DTT, NH4HCO3, TFA, urea) were from Sigma Aldrich (St. Louis, MO). Acetonitrile, HPLC grade, was from Fischer Scientific (Fair Lawn, NJ), and high purity water was prepared from DI water, in-house, by distillation.

### Cell culture

The cells were initially propagated in the manufacturer’s suggested growth media, i.e., SKBR3 in McCoy 5A, HMC3 in EMEM, HBEC5i in DMEM/F-12/ECGS, and NHA in ABM with SingleQuots growth supplements (FBS, EGF, insulin, L-glutamine, ascorbic acid, and gentamycin). After the third passage, 50 % of the culture media content was replaced with DMEM-HG, and after yet another 48 h the cell cultures were continued in 100 % DMEM-HG **(Figure 1A)**. All culture media were supplemented with FBS (10 %) and PenStrep (0.5 %) at all times, the astrocytes with the additional growth supplements, and the endothelial cells with ECGS. At ∼90-100 % confluence, the cell cultures were serum-starved for 24 h by growing them in DMEM-HG/PenStrep. Serum-free conditioned media (CM) collected from HMC3, HBEC5i and NHA were mixed and spun down by centrifugation at 500 x g for 5 min to remove the floating cells. Clarified CM was added to 24 h serum-starved SKBR3 cells which were grown for another 24 h in the presence of the secreted factors. The SKBR3 cells that were used as control were left under serum-starved conditions for a total of 48 h. Morphological differences between treated and non-treated SKBR3 cells were visualized by using phase-contrast and immunostaining microscopy and a Nikon Eclipse TE2000 U epi-fluorescence inverted microscope (Nikon Instruments Inc., Melville, NY) equipped with the NIS-Elements Advanced Research imaging platform. Mouse monoclonal anti-HER2 primary antibody (Santa Cruz sc-08) and mouse IgG-kappa binding proteins conjugated to CruzFluor488 were purchased from Santa Cruz Biotechnology (Dallas, TX). The cells were harvested by trypsinization and spun down to pellets which were flash frozen at -80 °C until further use. DNA content analysis for cell cycle stage assessment was performed by FACS using a FACSCalibur flow cytometer (BD Biosciences, San Jose, CA). Three independent biological replicates of control and treated cells were performed [control: 69-72 % in G1 (CV=9 %), 8-13 % in S, 2-7 % in G2; treatment: 70-90 % in G1 (CV=13 %), 11-14 % in S and 0.3-4 % in G2]

**Figure 1.**
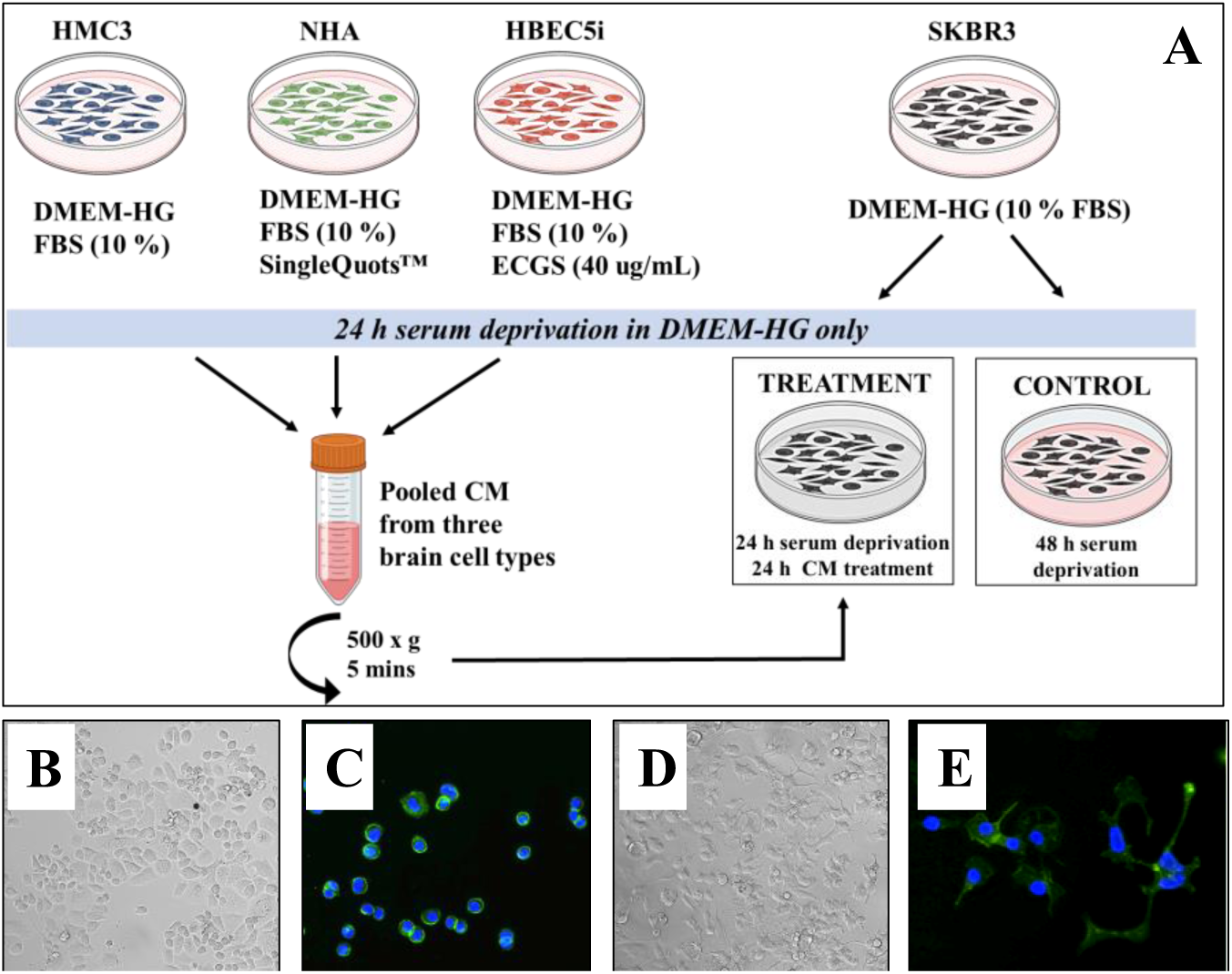
Overview of the experimental setup. **(A)** Schematic outline of the cell culture. Three biological replicates were established for HMC3, HBEC5i and NHA. All cells were grown in DMEM-HG medium under serum-deprived conditions for 24 h after which the media was collected and pooled from the three brain cell types to treat the SKBR3 cells for another 24 h. SKBR3-control cells were maintained in serum-free media for 48 h. **(B)** Phase contrast microscopy image of non-treated SKBR3 cells under 20X magnification. **(C)** Immunofluorescence microscopy of non-treated SKBR3 cells labeled with anti-HER2 antibody and detected using an anti-mouse IgG secondary antibody conjugated to CruzFluor488. **(D)** Phase contrast microscopy image of CM-treated SKBR3 cells under 20X magnification shows development of filopodia/lamellipodia type protrusions from the surface of the cells. **(E)** Immunofluorescence microscopy of treated SKBR3 cells labeled with anti-HER2 antibody and detected using an anti-mouse IgG secondary antibody conjugated to CruzFluor488.

### Cytokine array analysis

Protein profiling of the secreted factors from each cell culture was performed using the Proteome Profiler Human XL Cytokine array kit (R&D systems, Minneapolis, MN) according to the manufacturer’s recommended protocol. CM from HMC3, NHA and HBEC5i, at 90 % confluence, was collected after 24 h of serum starvation. CM from SKBR3 cells was collected after 24 h arrest + 24 h of stimulation with CM from brain cells (treatment), and after 48 h of serum starvation alone (control). After collection, the conditioned media were centrifuged at 500 x g for removal of floating cells, and 2 mL of the supernatant solution was added to the blocked membrane arrays and left to incubate overnight at 4 °C. After rinsing three times with wash buffer, the membranes were incubated with the antibody cocktail for 1 h at room temperature followed by a 30 min incubation with Streptavidin-HRP solution. The membranes were once again rinsed with the wash buffer and the signal was developed using the ChemiReagent mix provided in the kit. The chemiluminescent signal was captured using a ChemiDoc™ Imaging System (BioRad, Hercules, CA) and the raw images were analyzed for pixel intensities using the EMPIRIA Studio software (https://www.licor.com/bio/empiria-studio/). Background subtraction was performed by the software’s adaptive background subtraction method to minimize variability and improve the consistency of data from each array. The signal intensities from the duplicate cytokine spots on each array were averaged and normalized to the average of reference spots from all arrays that were exposed to the secretome of serum-starved cells. Two independent microarray measurements were acquired from two biological replicates of each cell line and further averaged for measuring differences in protein abundance. Only spots with averaged normalized pixel intensities >2000 were used for comparative purposes.

### Cellular fractionation into nuclear and cytoplasmic proteins

The cells were separated into nuclear and cytoplasmic fractions prior to MS analysis [20,21]. The isolation of nuclear proteins was achieved with the CelLytic™ NuCLEAR™ extraction kit (Sigma Aldrich, St. Louis, MO) using the manufacturer’s recommended protocol. In brief, the cells were disrupted under ice-cold conditions with a hypotonic buffer in the presence of protease inhibitors (Sigma product P8340), Na2VO3/NaF phosphatase inhibitors (1 mM), DTT (1 mM), and Igepal. The lysed cells were centrifuged at 4 °C for 1 min at 15,000 x g. The clarified supernatant containing the cytoplasmic components was set aside until further use. The nuclear pellet was further disrupted with a high-salt containing buffer, supplemented with the same inhibitors, by applying constant agitation using a vortex mixer for 30 min at 4 °C. The lysate was sonicated and centrifuged at 4 °C for 10 min at 15,000 x g to generate the nuclear protein fraction. The total protein concentration in the cell extracts was measured using the Bradford assay.

### Protein digestion and sample preparation for MS analysis

For MS analysis, the protein extracts were denatured for 1 h at 57 °C, pH∼8, in the presence of urea (8 M) and DTT (5 mM). The samples were diluted 10-fold with 50 mM NH4HCO3 prior to tryptic digestion. In-solution digestion was performed overnight on a shaker set at 37 °C using sequencing grade trypsin (Promega, Madison, WI) at ∼1:25 w/w enzyme/protein ratio. The reaction was quenched with TFA and the peptides were isolated and desalted using reverse phase C18 and SCX cartridges (Agilent technologies, Santa Clara, CA). Tryptic peptide mixtures were evaporated with speed vacuum and dissolved in 2 % acetonitrile solution acidified with 1 % TFA at a final concentration of 2 μg/uL.

### LC-MS analysis

The peptide samples (1-1.5 μL) were analyzed with a Q-Exactive hybrid quadrupole-Orbitrap mass spectrometer (Thermo Fisher Scientific) equipped with a nano-electrospray source operating at 2 kV at a capillary temperature of 320 °C and coupled to an EASY-nLC 1200 UHPLC system (Thermo Fisher Scientific) [21]. The nano-LC separation was performed on a 75 μm i.d. reverse-phase EASY-Spray column ES802A (250 mm long) packed with 2 μm C18/silica particles (Thermo Fisher Scientific). Column equilibration was performed with buffer A (H2O 96 %, CH3CN 4 %, TFA 0.01 %) and the peptides were eluted in a 2 h long gradient of buffer B (H2O 10 %, CH3CN 90 %, TFA 0.01 %) at a 250 nL/min flow rate. The MS was operated in a data-dependent acquisition mode with a full-scan range of 400-1600 m/z, resolution 70,000, AGC target 3E6, and maximum IT 100 ms. Tandem MS/HCD spectral acquisition was performed with a quadrupole isolation width of 2.4 m/z, 30 % normalized collision energy, resolution of 17,500, AGC target 1E5, maximum IT 50 ms, loop count of 20, and dynamic exclusion 10 s. To improve the consistency and reproducibility of MS measurements, each sample was injected three times. Relevant quantitative results were validated via parallel reaction monitoring (PRM) accomplished with quadrupole isolation width of 2 m/z, resolution of 35,000, AGC target (1-2)E5, and maximum IT of 110 ms.

### Database searching and protein identifications

Tandem mass spectra raw files were processed with the Proteome Discoverer 2.5 software package (Thermo Fisher Scientific, Waltham, MA), using the Sequest HT search engine, a minimally redundant/reviewed *Homo sapiens* database (UniProtKB, 2019 download), and two processing workflows that utilized either (a) the Percolator or (b) the Target/Decoy peptide spectrum match (PSM) validator nodes to discriminate between correct and incorrect PSMs and calculate the associated scores and statistical features (e.g., q-values, FDRs). The Sequest search parameters were set for fully tryptic precursor peptides with a maximum of 2 missed cleavages, min of 6 and max of 144 amino acids, 400-5000 m/z, dynamic modifications allowed on Met (+15.995 Da/oxidation) and on the protein N-terminal (+42.011 Da/acetylation), and precursor mass tolerance of 15 ppm and fragment ion tolerance of 0.02 Da. The Percolator node was run by using a concatenated Target/Decoy strategy and default parameters, i.e., q-value based validation, max delta Cn of 0.05, and strict/relaxed peptide FDRs of 0.01/0.05. Chromatographic peak identification across the different samples was performed by adding the Minora Feature Detector node in the processing workflow. Parameters for this node were set to min trace length of 5, S/N threshold of 1, max deltaRT 0.2, and PSM confidence set to at least high (FDR<0.01) for feature to ID mapping. The Target/Decoy PSM Validator node also used a concatenated Target/Decoy selection strategy, but the PSM and peptide strict/relaxed FDRs were set to 0.01/0.03. In the consensus workflows, the PSMs with Xcorr<1 were filtered out, the peptide group modification site probability threshold was 75, only rank 1 peptides were matched to parent proteins, and protein grouping was performed by enabling the strict parsimony principle. The Protein Validator nodes had the confidence thresholds set to FDRs of 0.01/0.05 when using the Percolator, and to 0.01/0.03 when using the Target/Decoy Validator nodes, respectively, in the two processing workflows.

### Label-free quantification and statistical analysis

Six datasets, i.e., three biological replicates of the treated and three of the non-treated cells, were considered for quantitation. Nuclear and cytoplasmic datasets were treated separately at the processing stage but combined later to enable a better understanding of the biological context. Protein abundances were calculated based on the summed peptide peak areas assigned to a particular protein, when using the Percolator node, and based on the count of correct PSMs, when using the Target/Decoy node. For peak area measurements, the chromatographic peaks were aligned with the Feature Mapper node, allowing for a max RT shift of 15 min and a peptide mass tolerance of 15 ppm. Global drifts in peptide abundance measurements were adjusted by normalizing the data based on total peptide abundance in a given sample (all peptides were used for normalization). Missing values were handled by enabling the imputation mode in the Precursor Ions Quantifier node with the low abundance resampling parameter (lower 5 % of detected values). Log2 fold changes (FC) of protein abundances in SKBR3 treated vs. non-treated datasets were calculated by using a pairwise approach, *i.e.,* by calculating the protein ratios as the median of all possible pairwise peptide ratios (unique and razor, excluding modified peptides). Statistically significant changes were derived by applying a *t-test* (background based). For PSM count-based abundance measurements, the datasets were normalized based on the average of total PSMs identified in each of the six datasets, one PSM was added to each protein to account for the missing values, and a *t-test* was performed for assessing significance. For both scenarios, only proteins that were matched by at least two unique peptide sequences, and for which the Log2(FC) was either ≥1 or ≤(-1) with p-value<0.05, were included in downstream analysis for biological interpretation. The PRM data were analyzed with Skyline (https://skyline.ms) [22], using mass spectral libraries created from the SKBR3 MS raw files. Peptides having a minimum of 5 transitions with a dot product (*dotp*) score >0.9 were considered for validation.

### Biological data interpretation

Biological relevance of the differentially expressed proteins, i.e., GO enrichment, pathway analysis, and visualization of proteins that displayed statistically significant differences in peak areas or PSM counts was performed by using publicly available bioinformatics tools provided by STRING, DAVID, KEGG, Reactome, GeneCards, and Cytoscape. Biological schematics were created with tools provided by Biorender.com.

## Results

To explore the early response of HER2+ breast cancer cells to the biological effects of the brain microenvironment, the SKBR3 cells were cultured using serum-deprived conditions in the presence (T-treated) and absence (C-control) of conditioned medium combined from serum-deprived astrocytes (NHA), microglia (HMC3) and brain endothelial (HBEC5i) cells **(Figure 1A)**. It was anticipated that the combined CM from all three brain cells would mimic the microenvironment of the metastasized tumor cells in the perivascular niche. The study was conducted in the absence of fetal bovine serum that is typically used for cell culture to avoid the interference of the broad range of components of undefined concentration present in the serum (e.g., growth factors, hormones, enzymes, proteins, carbohydrates, etc.), as well as the variable batch-to-batch composition that can affect the reproducibility of findings [23]. The cells were depleted of serum for 24 h prior to treatment to ensure that the impact of the treatment on SKBR3 cells stems only from the neural cell CM and not prior FBS supplementation. The control (C) and treated (T) nuclear (N) and cytoplasmic (C) cellular subfractions were annotated as follows: CN1-CN3 nuclear fractions of control cells, CC1-CC3 cytoplasmic fractions of control cells, TN1-TN3 nuclear fractions of treated cells, and TC1-TC3 cytoplasmic fractions of treated cells.

### Morphological characteristics of CM-treated SKBR3 cells

Under basal conditions, the SKBR3 cells displayed a grape-like morphology with a lower intercellular adherence (**Figure 1B**), as it has been confirmed by previous studies [24]. The cells stained strongly for the ERBB2 receptor which is known to be present in high abundance on the surface of the cells (**Figure 1C**). Upon treatment with the CM, we observed the development of protruding structures resembling lamellipodia and filopodia emanating from the surface of cells, which could be also visualized by ERBB2 staining (**Figure 1D, 1E**). DNA content analysis by flow cytometry revealed small but consistent differences, the treated SKBR3 cells displaying a larger proportion of cells in the G1 stage of the cell cycle in comparison to the controls [83 % (CV 13 %) treatment *vs* 74 % (CV 9 %) control], suggesting an inhibition of growth upon treatment with the CM. Nonetheless, the proteomic data (see further) revealed a more complex response to the CM treatment, that branched out into multiple biological processes beyond cell cycle arrest such as metabolism, adhesion, and immune response.

### Analysis of conditioned media using cytokine arrays

Membrane-based cytokine arrays were used to measure the relative concentration of 105 different components in the CM of serum-deprived brain and cancer cells and assess the factors relevant to intercellular communication that may affect the behavior of cells (**Figure 2**). Detailed results of the replicate cytokine measurements including images and raw and normalized pixel intensities are provided in **Supplemental file 1**. The functional impact of components that displayed noticeable pixel intensities (>∼2000) was evaluated. A set of ∼50 proteins of interest emerged from the analysis: chemokines and interleukins, growth factors, and various other factors with roles in growth/development, angiogenesis, adhesion, migration, and immune response. The group of proteins with increased abundance in at least one of the brain cell secretomes included pro-inflammatory chemokines (CCL2, CCL5, CCL7/MCP3, CXCL1, CXCL5, CXCL8/IL8, CXCL10, CXCL11), homeostatic chemokines (CXCL12/SDF1), pro- and inflammatory cytokines and factors (IL6, IL15, TNFSF13B, FLT3LG, RBP4, CD14, CSF1/2), pro-inflammatory inhibitors and homeostatic cytokines (IL18BP, PTX3), growth and cell activator factors (BDNF, IGFBP2/3, PDGFA, GDF15, TNFSF13B, ANGPT1/ANGPT2, CSF1/2, CHI3L1, SPP1, DKK1), and cell adhesion and extracellular matrix (ECM) remodeling molecules (ICAM1, VCAM1, MMP9, THBS1, SERPINE1, PLAUR) (**Figure 2A-C** and **2F**).

**Figure 2.**
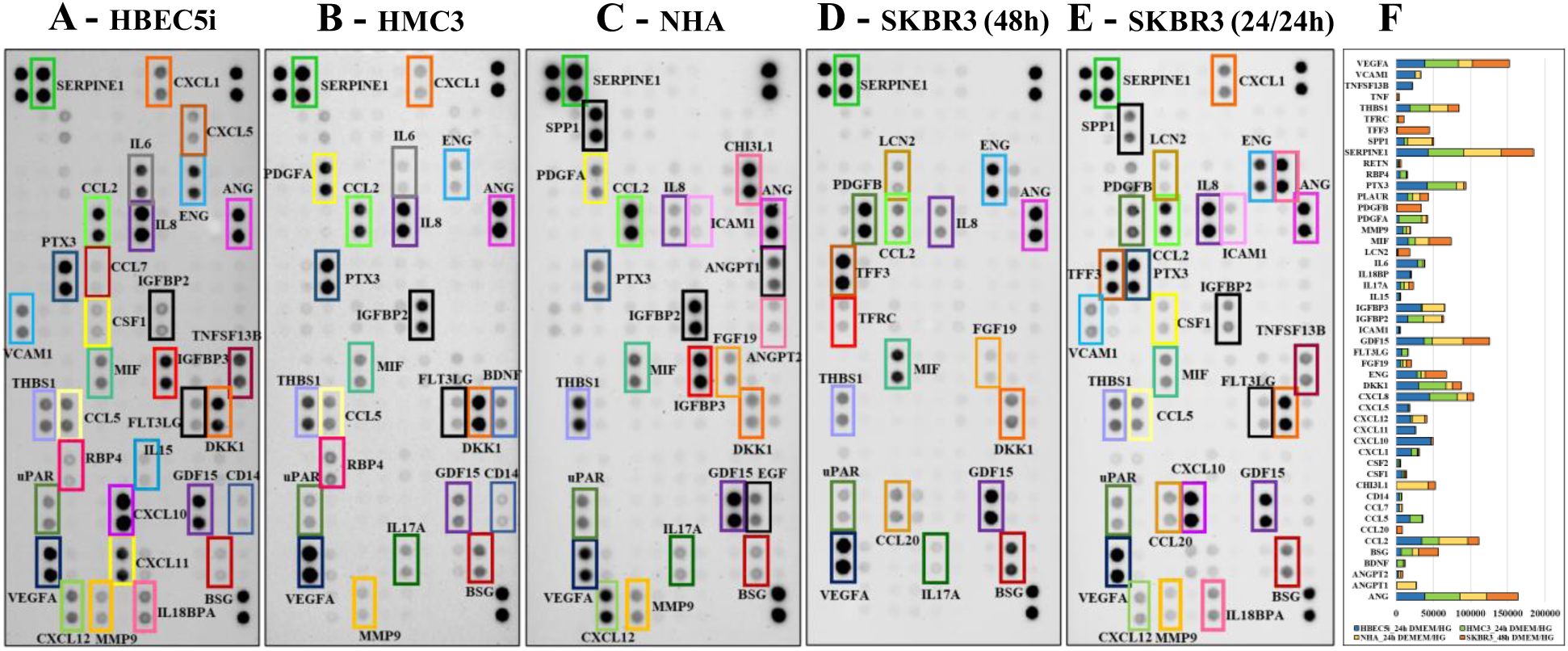
Results from the cytokine microarray profiling of conditioned medium collected from serum-deprived cell cultures of **(A)** HBEC5i, **(B)** HMC3, **(C)** NHA, **(D)** SKBR3-control, and **(E)** SKBR3-stimulated cells. **(F)** Bar chart showing the average pixel intensities of the set of cytokines identified in the CM collected from the serum-starved brain and SKBR3-control cells (pixel intensities >2000). Color code: blue-HBEC5i, green-HMC3, yellow-NHA, orange-SKBR3.

The group of proteins with increased abundance in the SKBR3 secretome (not necessarily unique to SKBR3) included cytokines (pro-inflammatory MIF, CCL20, IL17A, and anti-inflammatory TFF3) and several groups of factors and receptors with relevance to growth and development (PDGFB, FGF19, TFRC), and ECM remodeling and invasiveness (LCN2, BSG) (**Figure 2D** and **2F**). Angiogenesis factors such as VEGFA, ANG and ENG were secreted by all cells, with ENG being more abundant in the HBEC5i and SKBR3 secretomes. A schematic representation of a subgroup of cytokines that provided visually discernable spots (pixel intensity>7000), and of which some displayed discrepancies in abundance in the cell secretomes, is provided in a node/edge networked configuration in **Figure 3A** with edges indicating detectability in each of the four cell lines. A cutoff of ∼3-fold change (FC) in normalized pixel intensities in the group of HBEC5i/HMC3/NHA brain cells and SKBR3 was used to indicate unique association with a particular cell line.

**Figure 3.**
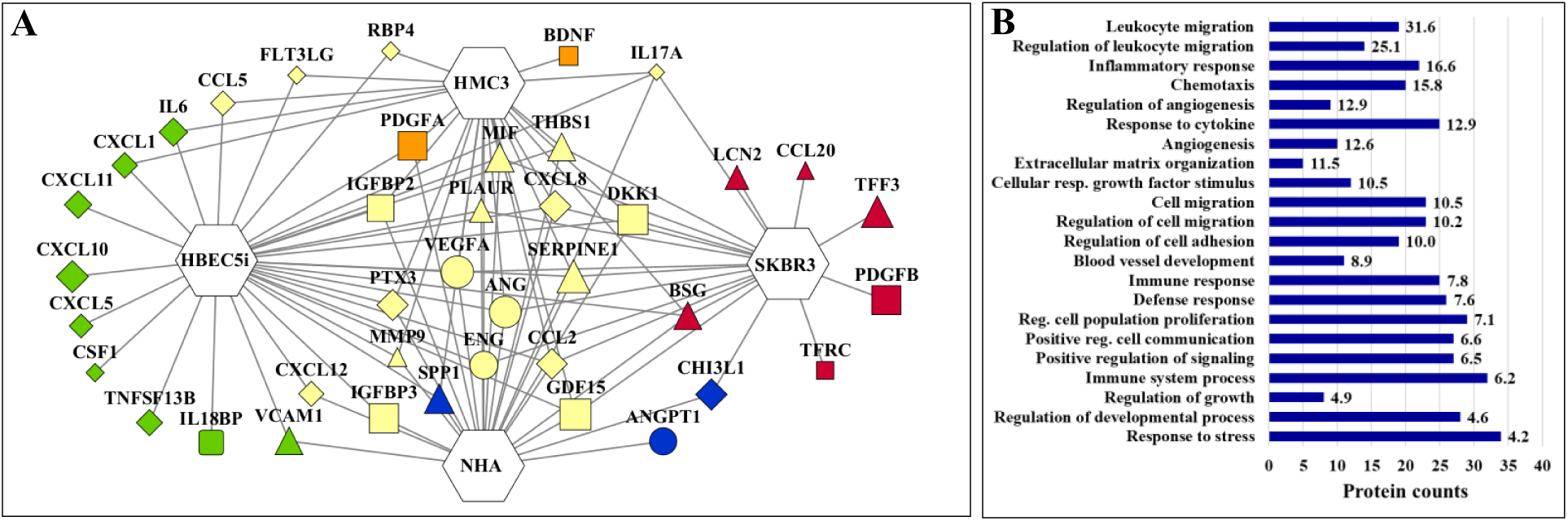
Overview of cytokines that displayed discrepancies in abundance between the brain and the SKBR3-control cell secretomes. **(A)** Diagram representing the detectability of cytokines in the serum-starved brain and SKBR3 cells (nodes represent gene names; edges connect the nodes to the cell lines in which the cytokines were detected; nodes and edges are shown only for nodes with a normalized pixel intensity greater than ∼7000). Symbol-code [size is proportional to log2(protein spot abundance)]: ▯ Growth/proliferation/survival/development/stress response, ◊ - Stimulation of immune responses and inflammatory conditions, Ο - Angiogenesis, Δ - ECM remodeling/adhesion/ migration/invasiveness. Color-code [indicates the cell line in which the microarray cytokine spot was most intense]: Red-SKBR3, Green-HBEC5i, Orange-HMC3, Blue-NHA, Yellow-cytokine secreted by multiple cell lines. **(B)** Bar chart of representative GO biological processes supported by all proteins with pixel intensity >2000 (FDR<5%). The labels associated with each biological process indicate fold enrichment.

#### Inflammatory cytokines

With few exceptions, all pro-inflammatory cytokines with elevated abundance in the brain cell secretomes were secreted at the basal level by the HBEC5i cells, with only a few being secreted in similar abundance by the HMC3 (CCL2, CCL5, CXCL8/IL-8, PTX3, RBP4, FLT3LG, CD14) and NHA (CCL7, CXCL8) cells. Owing to their prime location in the bloodstream, the endothelial cells are the first to come in direct contact with any infectious entity in the body. Therefore, these cells advanced mechanisms for the recognition of damage-associated molecular patterns (DAMPs) for inciting an immune response and for recruiting immune cells to the site of pathogenesis [25]. Many of these functions are facilitated by the inflammatory cytokines and chemokines that are released by the endothelial cells in their microenvironment. Elevated levels of TNFSF13B, a cytokine involved in the regulation of immune responses and stimulation of B-/T-cells, were observed only in the HBEC5i cell secretome. IL6 was also preponderantly expressed in the HBEC5i secretome. Along with its well-known inflammatory functions, this cytokine has been implicated in regulating the proliferation, angiogenesis, invasion, and metabolism of cancer cells. IL6 mediates these outcomes primarily via the STAT3 or NFKB signaling axis, and induces the expression of factors which promote enhanced malignancy (GM-CSF, CCL2, MMP, VEGF) [26]. Previous studies have confirmed a basal level expression of CCL2 and CXCL8/IL8 by the human brain endothelial cells, however, CXCL10 and CCL5 were shown to be expressed only upon pro-inflammatory cytokine stimulation [27]. The levels of CCL2 were relatively constant in all cell secretomes, but there is accumulating evidence that CCL2 induces angiogenesis via increased VEGFA expression, while CXCL5, CXCL8/IL8 and CXCL1 directly affect tumor growth and metastasis by enhancing blood vessel supply through neovascularization [28]. On the other hand, CXCL10 and CXCL11 exert angiostatic effects and initiate T-cell or NK-cell mediated immune responses [28]. Previous studies suggested that CCL7 could be involved in promoting tumor invasion and metastasis, but tumor suppressor effects of this cytokine have been also identified [29]. The detection of CCL7 in this work was, however, just above the intensity threshold setting in HBEC5i and NHA cells. IL15 was also observable in low abundance, and only in the secretome of HBEC5i cells. By stimulating the proliferation and activation of immune cells (NK, B- and T-cells), IL15 has many protective and anti-tumor roles. It is explored as a potential therapeutic agent by itself or as a target for the development of IL15 agonists and a number of immunotherapies [30]. IL18BPa (IL18 binding protein) which is an inhibitor of the proinflammatory IL18 cytokine was upregulated in HBEC5i, while the homeostatic/tissue remodeling PTX3 (Pentraxin-3) in the HMC3 and HBEC5i secretomes. IL18BPa is believed to play a buffering role for IL18 [31], and PTX3 to exert multiple tumor suppressive and supportive roles via complex involvement in mediating immune responses, ECM remodeling and angiogenic programs [32,33]. CXCL12 (SDF1α) was elevated in the HBEC5i secretome relative to SKBR3, and while this is a homeostatic chemokine, it is known to exert various roles in pathogenic conditions [34,35]. Different splice variants have specific activities, which are also controlled by PTMs [34], and hypoxia and growth arrest in different cell types can elevate the expression level of this cytokine [34]. Increased CXCL12 levels produced under hypoxia by ovarian cancers have been shown to promote the expansion of endothelial cells and angiogenesis [36]. Altogether, aberrant expression of CXCL12 in the tumor microenvironment was correlated with tumor growth, proliferation, inhibition of apoptosis, and driving cancer cell migration to distant sites [35]. The cytokine signals along the MAPK, PI3K/AKT, Wnt and NFKB pathways to induce the expression of VEGF, FGF, cyclooxygenase-2 and IL6, all of which are critical mediators of angiogenesis [35]. CXCL12 was also observed in the NHA medium. Additional pro-inflammatory proteins such as the Fms-Related Tyrosine Kinase 3 Ligand and growth factor (FLT3LG), retinol binding protein 4 (RBP4), and receptor CD14, *albeit* in lower abundance than the other cytokines, complemented the ability of HBEC5i cells to stimulate the proliferation or activity of immune cells [37]. In addition, the cell adhesion glycoproteins VCAM1 and ICAM1 with roles in mediating cell adhesion, motility and maintaining tissue architecture, were also predominantly expressed in the HBEC5i secretomes, especially VCAM1 (ICAM1 was observable in low abundance). VCAM1 is overexpressed on endothelial cells under inflammatory conditions to allow for the adhesion and rolling of immune cells on the endothelium surface [38]. It has particular relevance during cancer metastasis, as it is one of the molecules that tumor cells bind for transmigrating across the endothelial barrier [38].

In microglia, the presence of IL6, CXCL8/IL8, CCL5, PTX3 was consistent with the role of these cytokines in the regulation of leukocyte activation and chemotaxis that is invoked in the brain in the case of neuropathological and inflammatory conditions, and the detection of PTX3 and FLT3LG with the involvement of microglia in mediating inflammation and phagocytosis, respectively [39]. Likewise, the secretion of CCL7 by NHA cells was associated with promoting microglia-mediated inflammation after brain injury [40].

In the SKBR3 secretome, pro-inflammatory cytokines were present, but generally in low abundance (excepting IL17A and MIF which were also present in the brain cell secretomes) or had mostly tumor-promoting functions (CCL20). Cytokines such as IL11, IL22, RETN and TNF which are associated with inflammatory responses were observable at very low intensities, but their emerging role in promoting the progression of cancer has been recognized [41–44]. The other cytokines have been shown to promote cell renewal, tissue regeneration, recruitment of immune cells and activation of pro-survival and mitogenic signaling (IL17A [45]); induce cell invasiveness, angiogenesis and cell survival pathways (macrophage migration inhibitory factor (MIF [46]); or, stimulate invasiveness via MMP secretion and impart chemotherapeutic resistance (chemokine CCL20 [47]).

#### Growth factors

We further assessed the differences in the concentration of growth and stimulating factors between the SKBR3-control and the brain cells (**Figures 2A-D**) [37,48–51]. The analysis revealed that the level of many pro-tumor factors, especially of those that are implicated in growth/proliferation, angiogenesis, migration, and metastatic progression was either similar or higher in the SKBR3 than in the brain cell secretomes (PDGFB, FGF19, GDF15, MIF, LCN2, TFF3, VEGFA, ANG, ENG, ANGPT2, TFRC, SERPINE1, PLAUR, BSG) (**Figure 2F**, **Figure 3A**). Specific biological processes sustained by these proteins included regulation of ERK1/ERK2 and MAPK signaling cascades, cell adhesion, cell migration and invasion, blood vessel development and angiogenesis, and response to stress/defense and inflammation (**Figure 3B**). The multifunctional platelet derived growth factor subunit B (PDGFB) with roles in cell proliferation/survival and migration, in particular, was highly abundant in SKBR3 and essentially missing from all brain cells. GDF15 was expressed by all cells, and increased levels have been shown to be associated with advanced cancer and poor patient outcomes, as this factor promotes mechanisms of immune evasion in cancer cells and enhances cell viability and metastasis [48]. Others, however, have suggested a conflicting response to GDF15, when it was observed to induce apoptosis in some cancer cells [48]. Further evidence for the secretion of cancer progression supportive factors in SKBR3 was presented by the higher expression of lipocalin-2 (LCN2) which has important ramifications in EMT-related processes, invasion, and cell migration [49] and of the anti-inflammatory trefoil factor 3 (TFF3) which supports invasion and metastasis in several carcinomas [50].

All three brain cell types secreted some if not all of the above cancer-supportive factors. In a broader context, these proteins are involved in developmental processes, and regulate cell proliferation, mitogenesis, survival, tissue repair, and chemotaxis (**Figure 3B**). A few factors that were identifiable in SKBR3 were present in higher abundance in the brain cell secretomes (IGFBP2/3, PDGFA, SPP1, CSF1/2), some being particularly elevated in NHA (IGFBP2, ANGPT1, CHI3L1, SPP1) or HMC3 (PDGFA, BDNF) (**Figure 3A**). NHA exclusively expressed the angiopoietin (ANGPT1) and chitinase 3 like 1 (CHI3L1) proteins. Together with VEGFA, angiopoietins secreted by astrocytes are important for governing the angiogenic remodeling processes for the development of vasculature in the brain [52]. Increased levels of CHI3L1 have been associated with neurodegenerative and inflammatory brain diseases, but aberrant expression has been also observed in brain tumors and metastasis. It has been suggested that CHI3L1 expression by activated astrocytes may play a role in tumor progression, angiogenesis, immune escape, and resistance to therapeutic drugs [53]. It is also worth highlighting osteopontin (SPP1) which plays a crucial role in driving metastasis, invasion, angiogenesis, chemotaxis, and suppression of anti-tumor immune responses [54]. SPP1 is known to be secreted by both cancer cells and the cells of the TME, and to have an overall positive impact on the metastatic progression of cancer [54]. HMC3 revealed the expression of the BDNF involved in the development and survival of CNS cells. Interestingly, BDNF activates TrkB (Tropomyosin-Related Kinase B or Neurotrophic Receptor Tyrosine Kinase 2) and HER2 receptors and enhances the proliferation and survival of brain metastatic HER2+ breast cancer cells [55]. On the other hand, while the exact function of PDGFA in microglia is not fully understood, this growth factor is recognized for its important contributions to supporting cell proliferation, angiogenesis, migration, as well as chemotaxis [37]. The matrix remodeling proteins and enzymes (SERPINE1, PLAUR, PTX3, LCN2, BSG, THBS1, MMP9), together with the CAMs (ICAM, VCAM1) that were overexpressed in the HBEC5i secretome, formed a group that plays key roles in mediating cell-cell/cell-matrix interactions and ECM remodeling processes that ultimately affect, again, the adhesion, migration, and the invasion capabilities of cancer cells.

The culture medium collected from the SKBR3 cells grown in brain cell-conditioned media generated a signal for almost all factors that produced intense spots in the brain and SKBR3-control secretions (**Figure 2E)**. It was therefore difficult to conclusively determine which cytokines and biological processes were upregulated in the SKBR3 secretions after treatment with conditioned media from the brain cells.

### Proteomic analysis and quantification of the CM-treated vs non-treated SKBR3 cells

The impact of treating the SKBR3 cells with the brain cell-conditioned media was next evaluated quantitatively at the proteome level. The expected outcome was the identification of novel proteins that mediate an early response in SKBR3 cells to the factors that they encounter in a foreign microenvironment (i.e., the brain TME). Three biological replicates of SKBR3-treated (T) cells were compared to three biological replicates of SKBR3-control (C) cells. Two complementary approaches were used for assessing protein differential expression, i.e., based on LC peak area and PSM count measurements. The two approaches produced rather complementary results due to: (a) the actual use of two distinct methods for assessing protein abundance; (b) the use of two distinct approaches to validate the PSMs, with the area-based measurements being enabled by a workflow that included the Percolator node that uses a semi-supervised machine learning approach to distinguish the correct from the incorrect PSMs, in contrast to the PSM count measurements that relied on using the results from the Target and Decoy database searches; and (c) the possible software-related misrepresentation of area measurements because some chromatographic peaks did not have actual PSMs associated with them in all treatment and control samples. The complementarity of such results is not surprising when only few proteins change abundance or the changes in abundance are small, and it was noted in previous studies, as well [56]. Longer exposure of the SKBR3 cells (>24h) to the brain CM could have amplified the changes in the proteome profiles, however, the prolonged lack of serum from the culture medium would have biased the results by inducing an excessive stress response.

On the average, ∼3,200/4,700 proteins per cell fraction were identified in the SKBR3 cells when using the Percolator and Target/Decoy validator nodes, respectively, with over 70 % of proteins being identified with two or more unique peptides (**Supplemental file 2**). The reproducibility of protein identifications in biological replicates for the workflow that used the Percolator node is illustrated in **Figure 4** (# protein IDs, overlaps, PSM correlations), and the results of the quantitative comparisons in **Figures 5A-E** and **5G** [box and whisker plots of raw and normalized Log10(Protein Abundance) and Volcano plots] and **Supplemental file 3**. There were no major overlaps between the proteins that displayed differences in counts in the nuclear and cytoplasmic fractions, demonstrating the complementarity of the data (**Figures 5F** and **5H**). Taken altogether, the area and PSM-based upregulated datasets comprised 96/91 proteins, respectively, and the downregulated ones 256/46 proteins, combined from the analysis of both nuclear and cytoplasmic samples. A few representative proteins are illustrated in **Figure 6A**. Selected proteins were validated using PRM-MS analysis (**Supplemental file 4**).

**Figure 4.**
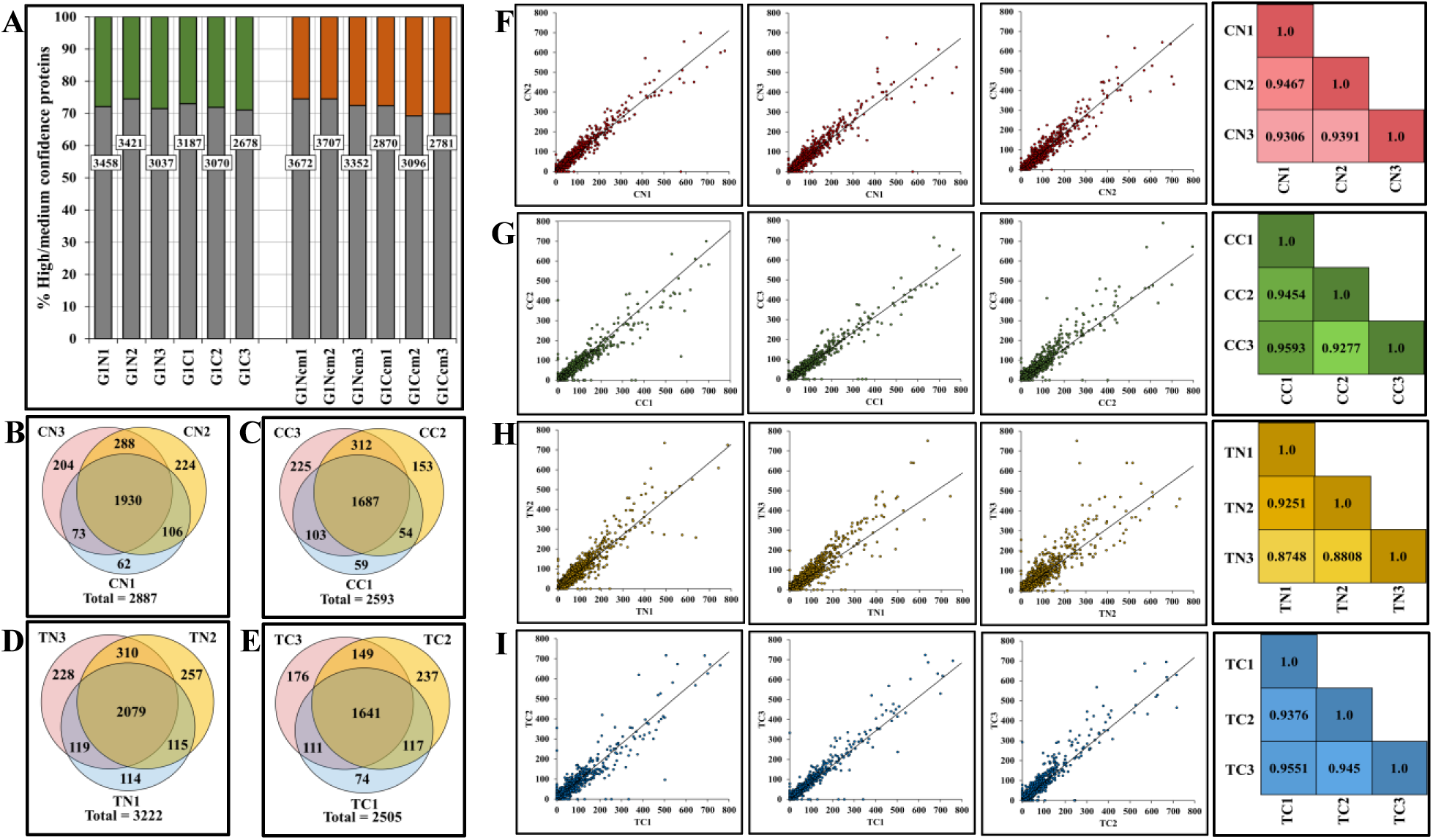
Evaluation of reproducibility in protein identifications. **(A)** Stacked bar chart showing the percentage of proteins identified by 2 or more unique peptides in a cell state and fraction (∼70 %, bars shaded in gray). The white boxes in the center display the total number of identified proteins in each cell state and fraction. (**B-E**) Venn diagrams displaying the overlap between the proteins identified with high/medium FDR and ≥2 unique peptides in three biological replicates of various SKBR3 cell fractions: **(B)** CN, **(C)** CC, **(D)** TN, and **(E)** TC. (**F-I**) Scatterplots displaying the degree of correlation between the PSMs of two biological replicates for **(F)** CN, **(G)** CC, **(H)** TN, and **(I)** TC datasets. Heatmaps on the right show the Pearson correlation coefficients for every comparison.

**Figure 5.**
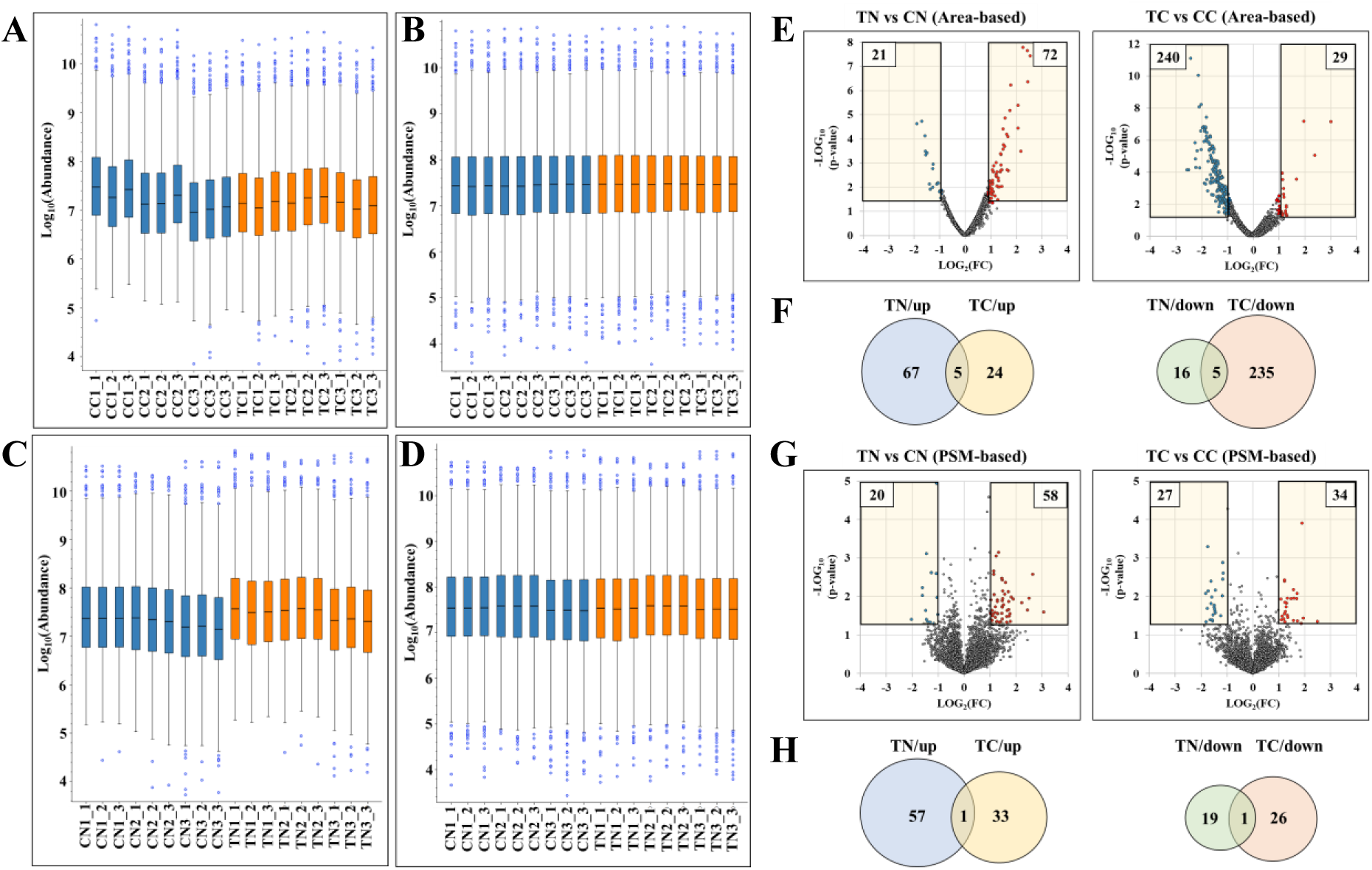
Overview of the proteomic label-free differential expression analysis approach. **(A-D)** Boxplots of area-based protein abundances for three biological and three technical replicates of SKBR3 cell fractions: **(A)** before and **(B)** after normalization for the quantitative analysis of TC vs CC fractions; **(C)** before and **(D)** after normalization for the quantitative analysis of TN vs CN fractions. **(E**, **G**) Volcano plots illustrating the results of differential expression analysis for proteins that displayed minimum 2-FC up- (red) or downregulation (blue) with a p-value ≤0.05 in TN vs CN and TC vs CC: **(E)** area-based measurements, and **(G)** PSM count-based measurements. **(F, H)** Venn diagrams showing the overlap between the proteins in the N and C fractions that were either up- or down-regulated in the area-based and PSM count-based measurements.

**Figure 6.**
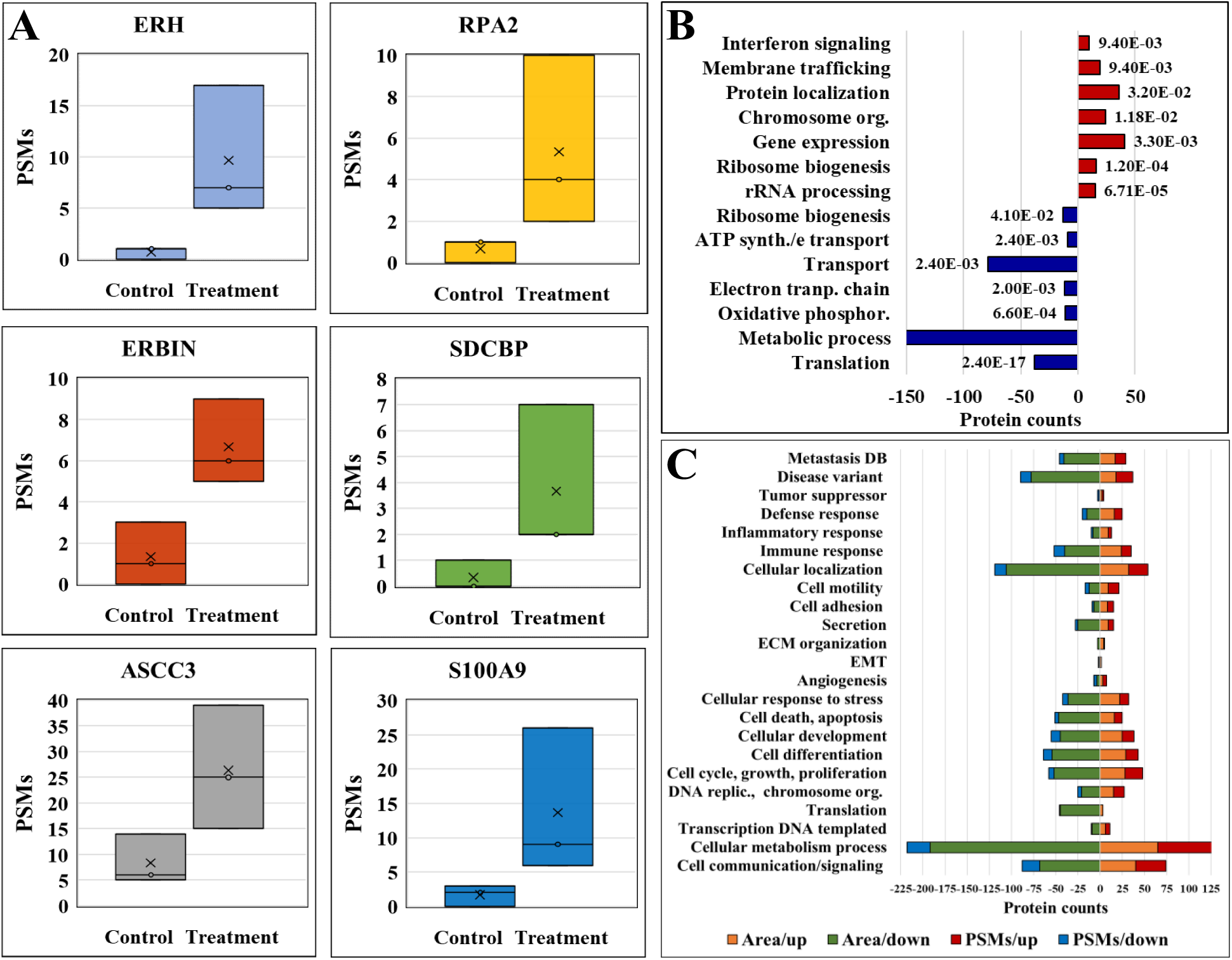
GO Biological processes represented by the proteins that changed abundance in the proteomic profiling of SKBR3 cells treated with CM medium from brain cells. **(A)** Box-whisker plots displaying the relative spectral counts for the selected proteins identified in the control and treated proteomic datasets. (**B**) Enriched biological processes represented by the combined nuclear/cytoplasmic proteins detected by the area and PSM-count measurements with increased or decreased abundance, respectively. **(C)** Protein counts that emerged from the area and PSM measurements mapped to GO biological processes of relevance to cancer (note: the proteins were mapped to biological processes by using GO controlled vocabulary terms).

Quantification was performed separately for the nuclear and cytoplasmic fractions, however, the results were combined for a thorough understanding of the cellular biology. The short lists of differentially expressed proteins yielded only very few enriched biological functional categories. When combined, however, the nuclear and cytoplasmic results, as assessed either via area or PSM measurements, revealed the main processes that were affected by treating the cells with brain cell conditioned medium. The processes that were sustained by proteins with increased counts were dominated by altered gene expression, chromosome organization, trafficking, protein localization, and interferon signaling. The processes that were supported by proteins with decreased counts (represented mostly by the cytoplasmic cluster generated by area measurements) were broadly associated with various metabolic and transport processes, and included an over-representation of cytoplasmic translation and electron transport/mitochondrial respiratory processes (**Figure 6B** and **Supplemental file 5**). Ribosome biogenesis was represented by proteins with both increased and decreased counts. Many of the above processes are mediated via posttranslational activation/deactivation or protein shuttling between various cellular organelles. Therefore, increased or decreased protein abundance measurements may not be reflective of actual protein up- or down-regulation, but rather of changes in PTMs or cellular location. Altogether, however, the results suggest that the treatment of cells with conditioned medium altered the cellular transcriptional/translational machinery with a net outcome of slowed metabolic processes. Ribosome biogenesis, for example, is an energy-intensive process, and a disruption in the availability of nutrients or energy in the cell that can affect any step of the biogenesis process leading to either up- or down-regulated ribosome components, has been shown to result in altered cell cycle progression or even cell death [57].

It is important to reiterate that upon treatment with the brain cell-conditioned media the SKBR3 cells were exposed to many inflammatory molecules that were either absent or present in low abundance in the secretome of the SKBR3-control cells. In the same time, the SKBR3-control cells secreted larger amounts of several important growth factors that may or may not have been at optimum levels in the brain cell-conditioned media to elicit a signaling response in the SKBR3 cells. Therefore, to derive a better understanding of the processes that were dysregulated in the SKBR3 treated cells, the proteins were further mapped directly to specific GO terms reflective of cancer-supportive processes and metastatic progression (**Figure 6C**). The complex interplay between these proteins in affecting the cell fate is exemplified in a Sankey diagram that presents a selected list of SKBR3 proteins with changed abundance, matched by at least three unique peptides, mapped to such processes (**Figure 7** and **Supplemental file 6).**

**Figure 7.**
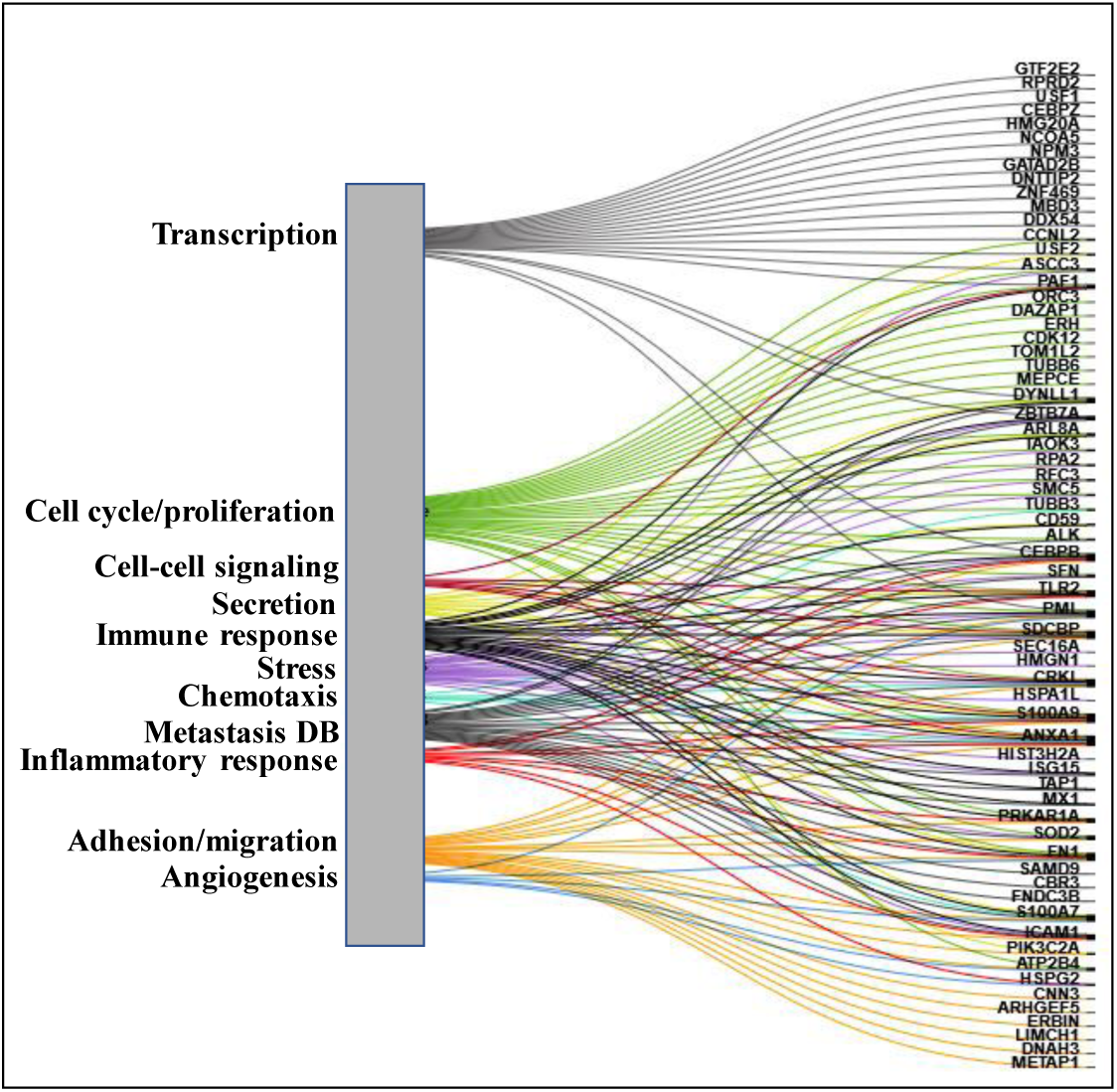
Sankey diagram illustrating the mapping of dysregulated nuclear/cytoplasmic proteins with increased abundance to key biological processes of relevance to metastatic progression of cancer (note: only proteins identified by at least three unique peptides are shown).

Based on our previous work on proteomic profiling of SKBR3 cells that has identified a broad range of receptors in the cell membrane (EGFR, FGFR, CSFR, IL, CD44/Cd47, IL, PDGFRA, TNFR, VEGFA receptor FLT1, ephrin, integrin, neuropilin NRP1, activin, chemokine ACKR2, adiponectin ADIPOR1, sortilin SORT1, low density lipoprotein LRP, syndecan SDC4) [58], a depiction of the possible interactions between the SKBR3 breast cancer and the brain cells is provided in **Figure 8**. Relevant SKBR3 receptors, along with the most abundant cytokines and growth factors secreted by the brain cells, are listed in the figure. The observed changes in the SKBR3 proteome were further interpreted based on the consideration that the SKBR3 cells that were exposed to the CM from brain cells experienced an environment deprived of optimal levels of growth factors needed for proliferation, but more abundant in inflammatory molecules. Proteins with similar trends in abundance changes, as emerged from the two quantification approaches, are discussed below.

**Figure 8.**
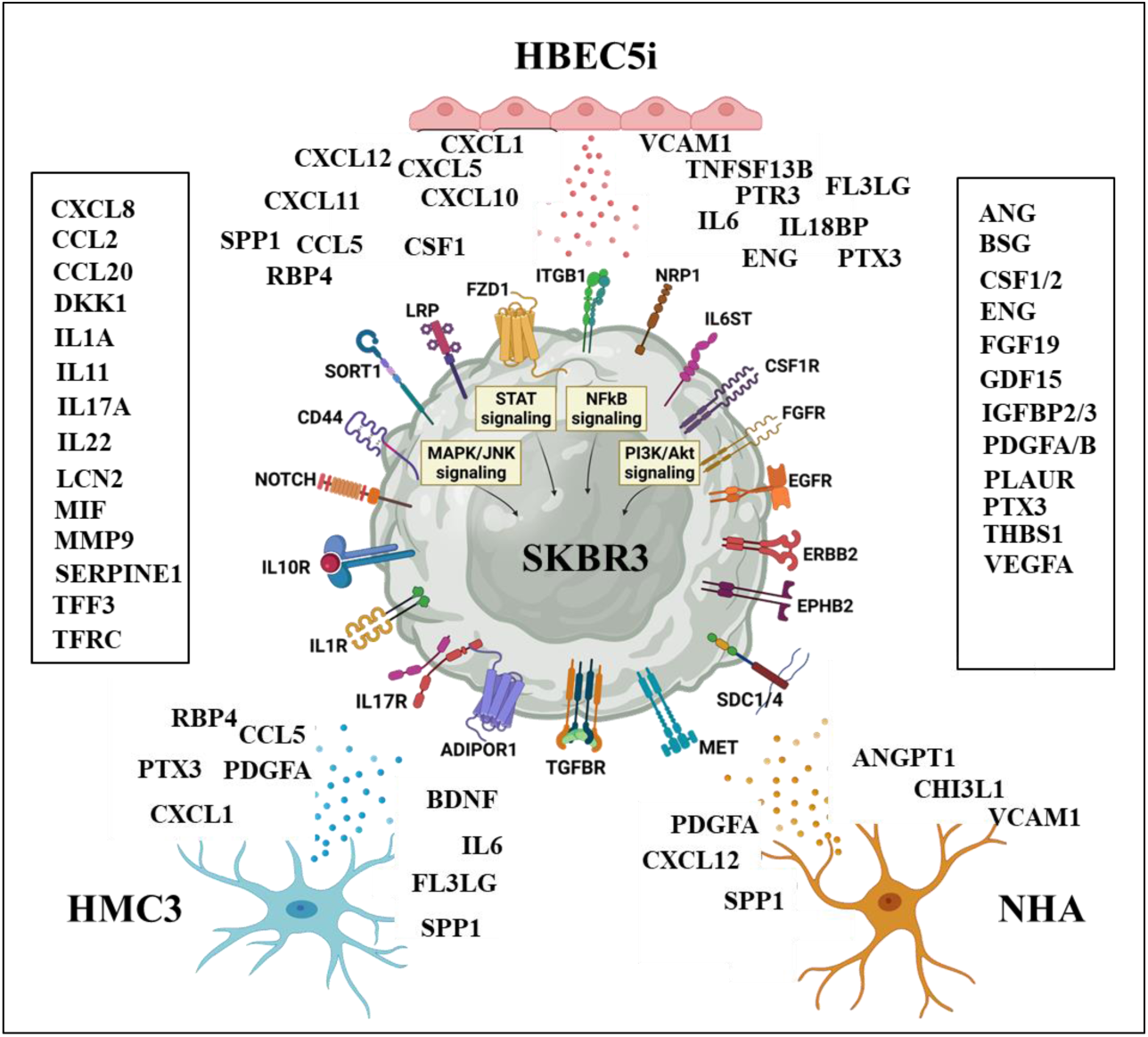
Illustration of receptors and secreted factors that can enable interactions between the SKBR3 breast cancer and brain cells. Relevant SKBR3 receptors were detected by proteomic profiling [58]. Abundant cytokines and growth factors secreted by brain cells, but missing or detected in low abundance in the SKBR3 secretome, are depicted near each cell line. Abundant cytokines and growth factors secreted in common by all brain and/or cancer cells are shown inside the sidebars.

## Discussion

### Effect of CM-treatment on cell cycle and proliferation

An interesting behavior of metastatic cancer cells is dormancy, a mechanism through which the metastatic cells temporarily exit the cell cycle to enter a state of quiescence and growth arrest, which allows them to survive the adverse conditions of a foreign niche [59]. Dormancy is activated in cancer cells to cope with environmental stress triggered by growth factor deprivation, oxidative stress, lack of ECM attachment and stromal cell interactions [59,60]. As a result, the quiescent cells significantly reduce their metabolic activity including biosynthesis of macromolecules, carbohydrate and lipid metabolism, and mitochondrial respiration [61]. A recent study demonstrated an enrichment of stress response and cell cycle related processes in the transcriptomics data derived from the dormant cancer cells of mouse and human origin [60]. In this study, a number of transcription factors and regulators were affected by the SKBR3 stimulation process with both initiator/activator (GTF2E2, USF1, CDK5RAP3, PML, CEBPB) and repressor roles (GATAD2B, PML, ZBTB7A).

In particular, ZBTB7A (zinc finger and BTB domain containing 7A protein), which is a transcriptional repressor of genes implicated in cell cycle progression and known to exert multiple roles in cancer cells that affect NFKB and Notch signaling, glycolysis, OXPHOS, transcription, chromatin organization, and DNA damage [62], warrants further investigation to provide clarity about its contribution to tumorigenesis. In contrast, CEBPB, a transcription factor involved in the regulation of a number of immune and inflammatory responses (see also later in the text), has been recognized for both its proliferative and anti-proliferative activities [37]. Clarifying the mechanisms of its functionality, however, is challenging, because the function of CEBPB in cancer cells is context specific, with diverse outcomes in different types of cancers, and because its activity is regulated by a number of different PTMs [63]. While on one hand CEBPB inactivates p53 expression to improve rates of survival and reduce apoptosis in cancer cells, it has also been shown to induce growth arrest in several types of cancers [63]. Several sets of proteins from the dataset also pointed to the possible alteration of signaling pathways that can modulate in concert cell cycle, proliferation, stress and inflammation, such as EGFR (ERBIN), p38/MAPK (TAOK3), PI3K (PI3KR2), DNA repair (RPA2, ASCC), TGFB (ZBTB7A, SDCBP), NFKB-mediated transcription and host-virus interactions (CDK5RAP3/LZAP), and anti-inflammatory processes (GNG5). TAOK3 regulates many important stress-responses in cells including apoptosis, metabolism, cell cycle, proliferation, cytoskeleton rearrangement, and inflammation [64]. Recently, it was found to induce drug resistance in breast cancer cells via NFKB activation and pathways that can have diverse outcomes, one of which is blockade of cell cycle progression and development of quiescence or dormancy [65,66]. Notable were also the upregulated replication protein A2 (RPA2) that responds to cellular stress by repairing damaged DNA at cell cycle checkpoints [37], and CDK5RAP3 and PML, due to their involvement in numerous cellular processes such as transcription regulation, DNA repair, growth suppression, apoptosis, and viral defense mechanisms. The last two proteins have been additionally recognized for their potential role in tumor suppression. Altogether, the results suggest that the cells demonstrated a complex response, rather than just a more pronounced cell cycle arrest. The cells possibly initiated adaptive mechanisms to survive the nutrient-limited stress conditions. In fact, cancer cell quiescence has also been linked to increased resistance to apoptosis [67]. These results further indicate that although the cells activated biological mechanisms in response to nutritional and inflammatory stress, the cells were not apoptotic.

### Effect of CM-treatment on cancer cell metabolism and response to stress

Scarcity of nutrients and unfavorable growth conditions drive cancer cells towards quiescence, which results in reduced metabolic activity, biosynthetic rates and respiration [61]. Recent studies have shown that dormant/quiescent cells switch their carbohydrate metabolism from mitochondrial OXPHOS to fatty acid oxidation, in order to secure a minimal supply of energy and prevent oxidative damage caused by the reactive oxygen species released during mitochondrial respiration [68,69]. Several metabolism associated proteins were downregulated according to the area-based measurements, including the ribosomal subunit proteins (RPLs, RPSs) involved in translation, the large neutral amino acid transporter (SLC7A5/LAT1), and the mitochondrial OXPHOS/electron transport and ATP production subunit proteins. Ribosome biogenesis is an energy consuming process, necessary for rapid cell growth and division, as a result of which ribosomes have a central role in a number of oncogenic signaling networks [57,70]. Under conditions of nutritional stress, cells can curtail the biogenesis of ribosomes to conserve energy, and, consequently, limit the G1/S progression of cells [71]. Reduced levels of solute carriers such as of the nucleotide-sugar transmembrane transporter (SLC35A4), that are implicated in cellular response to stress by regulating translation processes, corroborated the results. Lower expression of the large neutral amino acid transporter (LAT1/SLC7A5) was also suggestive of impeded proliferation in the CM-treated cells. The LAT1 transporter allows for an increased uptake of essential amino acids from the external microenvironment, in exchange for glutamine, which is pumped out of the cells [72]. LAT1 overexpression was found associated with highly proliferating cancer cells growing under low nutrient conditions and hypoxia, and is being evaluated for its therapeutic target potential [72]. The cluster also included proteins that mediate transport across the mitochondrial membrane, such as the glutamate carrier SLC25A22 and amino acid transporter SFXN1, all of which pointed toward restricted mitochondrial metabolism. These results correlated with the upregulation of the transcriptional repressor of genes involved in OXPHOS and glycolysis (ZBTB7A). Overall, the results indicate that the nutritional stress conditions induced by CM-stimulation impeded ribosomal protein synthesis pathways and reduced mitochondrial respiration. Furthermore, the inflammatory conditions mediated by the cytokines and chemokines triggered an overexpression of proteins that respond to these stimuli by regulating changes in cancer cell proliferation, growth and survival.

### Effect of CM-treatment on cancer cell adhesion/migration and EMT

Epithelial cells are characterized as having an apical-basal polarity, with junction proteins binding the cells together in a monolayer, attached to the basement membrane [73]. During cancer progression, tumor cells undergo EMT (epithelial to mesenchymal transition) which is marked by the loss of certain adhesion and junction proteins and acquisition of mesenchymal features, such as cytoskeletal rearrangements, that induce invasive and migratory capacities in the cancer cells [73]. The CM treatment of SKBR3 resulted in changes in the expression level of several proteins that have known implications in adhesion, migration and EMT-related processes.

Evidence for EMT type behavior and development of cell motility in the CM-treated SKBR3 cells was provided by the overexpression of syntenin-1 and annexin proteins. Syntenin-1 (SDCBP) activates cancer cell invasion and migration through actin remodeling processes during the late stages of cancer metastasis [74]. It colocalizes with growth factor receptors, the focal adhesion kinase complex, and other integrin-associated signal transducers to activate p38MAPK/JNK downstream signaling [74,75]. This, ultimately triggers cytoskeleton rearrangement processes, which regulate adhesion and migration in cancer cells [75]. A critical outcome of this pathway is the activation of mesenchymal proteins such as SLUG that induce EMT [75,76]. The cytoplasmic fractions of the treated cells displayed an upregulation of annexin A1 (ANXA1), which has the ability to bind the actin cytoskeleton, which in turn is crucial for the initiation of EMT, migration, and invasive properties in cancer cells [77]. The multifunctional ZBTB7A, as noted above, has been shown to also mediate EMT via the NFKB signaling pathway in a number of breast cancer cell lines [78].

A few additional proteins of potential interest included the downregulated YBX1 and LAMP1, and upregulated ICAM1 and S100 Ca-binding proteins. Notably, the DNA/RNA binding protein YBX1 participates in a range of cellular transcription/translation processes and augments metastatic processes such as invasion and cell migration [79], development of drug resistance [80], and elevated glycolysis [81]. Its activity, however, depends on phosphorylation, and further investigation is required to provide clarity on its activation status in the CM-treated SKBR3 cells. The lysosome associated membrane glycoprotein LAMP1 provides glycan ligands to selectins and promotes cell-cell adhesion in metastatic cancer cells [82]. ICAM1, an intercellular cell adhesion molecule, is involved in cell-cell and cell-ECM adhesion, and was found to be expressed by metastasized cancer cells as they interact with the endothelial cells to undergo trans-endothelial migration. S100 calcium-binding proteins bind to a variety of cell surface receptors on other cells to induce metastatic progression [83]. These proteins carry out, however, a broader range of functions that present relevance to metastatic processes, both in the intra- and extracellular environments. The intracellular functions of S100 proteins include involvement in various aspects of cell cycle regulation, promotion of cell survival and invasion via the NFKB signaling pathway, metabolism, and trafficking. The extracellular functions include involvement in pro-inflammatory, anti-microbial and death-inducing activities [37]. Binding to receptors on macrophages/microglia or endothelial cells facilitates their recruitment and angiogenesis at the site of tumorigenesis [83]. In this study, S100A7 and S100A9 were found upregulated in the CM-treated cells, while other isoforms (A4, A6, A13, and A16) appeared to be rather downregulated, pinpointing again to the need for careful interpretation of results and further studies.

### Effect of CM-treatment on the mechanisms of immune evasion

Upregulated ISG15 (interferon-induced ubiquitin-like protein) and MX1 (interferon-inducible protein p78 or myxoma resistance protein 1) were two proteins that were potentially reflective of the SKBR3 cell response to exposure to the cytokines from the conditioned medium. ISG15 plays multiple roles in innate immune responses that include antiviral activities, cell-cell signaling, and chemotactic activity [37]. It can act either in free form or by binding to target proteins, including MX1, a protein that impedes viral replication. Previous studies have demonstrated that ISG15 enhances EGFR recycling on the surface of breast cancer cells, as a result of which the cells display an increased membrane localization of EGFR and aberrant tumorigenic signaling [84]. A recent study identified MX1 as a potent inducer of migration, EMT, and invasion in cancer cells due to its association with EGFR signaling [85]. According to the cytokine arrays, a small increase in the abundance of interferon γ in the secretome of CM-treated SKBR3 cells was observable, but further studies would be needed to clarify the complex mechanisms that are responsible for the upregulation of these proteins. In the same time, the lysosomal thiol reductase IFI30 which is involved in MHC class I/II antigen presentation and which is known to be induced by interferon-γ in non-professional antigen presenting cells, was downregulated, suggesting a loss of MHC expression by the SKBR3 cells, which is a well-known mechanism used by cancer cells to evade immune cell recognition [86].

## Conclusions

The findings of this study provide preliminary, yet novel, insights into the early response of breast cancer cells to *in vitro* conditions that mimic the neural niche. The results led to the hypothesis that the unfavorable environmental conditions, possibly mediated by inflammatory molecules or suboptimal concentrations of growth factors, drive the cancer cells toward quiescence, and possibly dormancy, in a foreign metastatic niche. The pro-inflammatory chemokines and cytokines secreted by the brain cells have pleiotropic roles with anti-tumor activities and simultaneous functions in facilitating the recruitment of immune cells and promoting neovascularization to enhance blood vessel supply in a hypoxic TME. The pro-tumorigenic growth factors, on the other hand, can counteract the detrimental effects of the pro-inflammatory mediators and promote viability of cancer cells. Altogether, however, the SKBR3 cells secreted cancer-promoting growth factors in higher abundance than the brain cells, and displayed signs of impaired growth when subjected to the brain cell-conditioned medium. Such pilot studies can provide crucial insights into the initial response of cancer cells in a foreign niche, especially as the concept of dormancy is difficult to replicate experimentally under both *in vivo* and *in vitro* conditions. Furthermore, such exploratory results can pave the way for future research into critical questions such as how do cancer cells adapt and/or mutate to nutritionally deprived conditions, and how the combination of factors in the neural niche protects and later supports the revival of dormant cells. Finally, the extrapolation of this study to primary cells, fluids or tumors collected from brain metastatic patients can provide a stepping stone in the discovery efforts of novel drug targets and therapeutic strategies that target not just the cancer cells but also the tumor microenvironment.

## Data availability

The mass spectrometry raw files were deposited to the ProteomeXchange Consortium via the PRIDE [87] partner repository with the following dataset identifier: PXD046330.

## Author contributions

SA performed the experiments; SA and IML analyzed the data and wrote the manuscript; IML coordinated the work. All authors reviewed and approved the final version of the manuscript.

## Acknowledgment

This work was supported by an award from the National Institute of General Medical Sciences (Grant No. 1R01GM121920) to IML.

