## Supplemental file 2 for "Proteomic Insights into Breast Cancer Response to Brain Cell-Secreted Factors"

Summary of the total protein counts identified in the three biological replicates of the control and treated SKBR3 datasets.

(A) Workflow that included the Percolator validator node (PSM FDRs=0.01; only high confidence PSMs included in the results)

| Sample name | Treatment | # High confidence proteins (FDR=0.01) | # High confidence proteins with ≥2 peptides (FDR=0.01) |
|---|---|---|---|
| **CN1** | 48 h in serum-free DMEM-HG | 3458 | 2495 |
| **CN2** | | 3421 | 2548 |
| **CN3** | | 3037 | 2171 |
| **TN1** | 24h in serum-free DMEM-HG / 24h in CM from brain cells | 3672 | 2736 |
| **TN2** | | 3707 | 2761 |
| **TN3** | | 3352 | 2427 |
| **CC1** | 48 h in serum-free DMEM-HG | 3187 | 2327 |
| **CC2** | | 3070 | 2206 |
| **CC3** | | 2678 | 1903 |
| **TC1** | 24h in serum-free DMEM-HG / 24h in CM from brain cells | 2870 | 2077 |
| **TC2** | | 3096 | 2144 |
| **TC3** | | 2781 | 1943 |

(B) Workflow that included the Target/Decoy validator node (PSM FDRs =0.01/0.03; only medium and high confidence PSMs included in the results)

| Sample name | Treatment | # High/medium confidence proteins FDR=0.01/0.03 | # High/medium proteins with ≥2 peptides FDR=0.01/0.03 |
|---|---|---|---|
| **CN1** | 48 h in serum-free DMEM-HG | 4992 | 3982 |
| **CN2** | | 5133 | 4029 |
| **CN3** | | 4469 | 3616 |
| **TN1** | 24h in serum-free DMEM-HG / 24h in CM from brain cells | 5446 | 4173 |
| **TN2** | | 5469 | 4218 |
| **TN3** | | 4827 | 3844 |
| **CC1** | 48 h in serum-free DMEM-HG | 4745 | 3452 |
| **CC2** | | 4580 | 3372 |
| **CC3** | | 4012 | 3033 |
| **TC1** | 24h in serum-free DMEM-HG / 24h in CM from brain cells | 4306 | 3223 |
| **TC2** | | 4305 | 3243 |
| **TC3** | | 4263 | 3185 |
