## Supplemental file 4 for "Proteomic Insights into Breast Cancer Response to Brain Cell-Secreted Factors"

**Supplemental file 4**
**PRM-MS validation for selected proteins with increased counts in the CM-treated SKBR3 cells**

**CD59 (CD59 glycoprotein)**
**FEHCNFNDVTTR, Charge +2, m/z 741.82448**

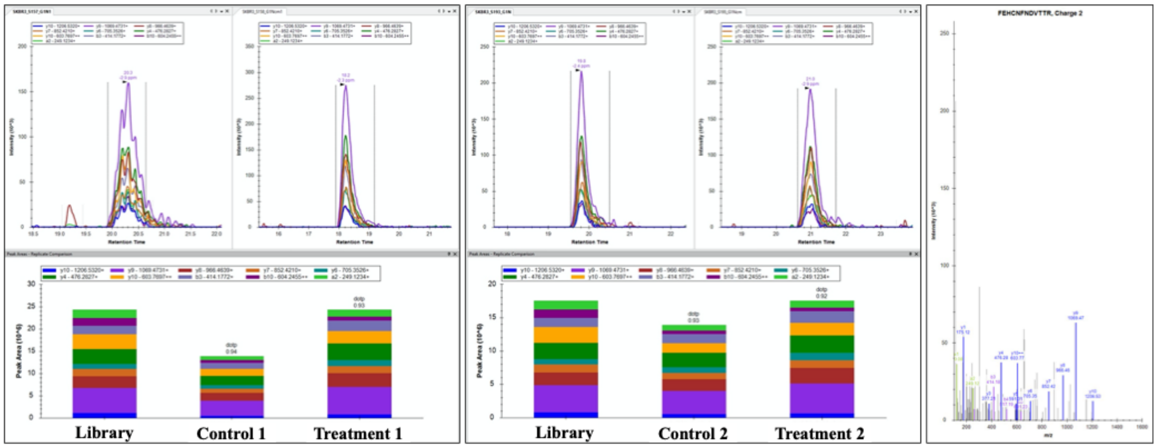


**ICAM1 (Intercellular adhesion molecule 1)**
**REPAVGEPAEVTTTVLVR, Charge +2, m/z 642.02187**

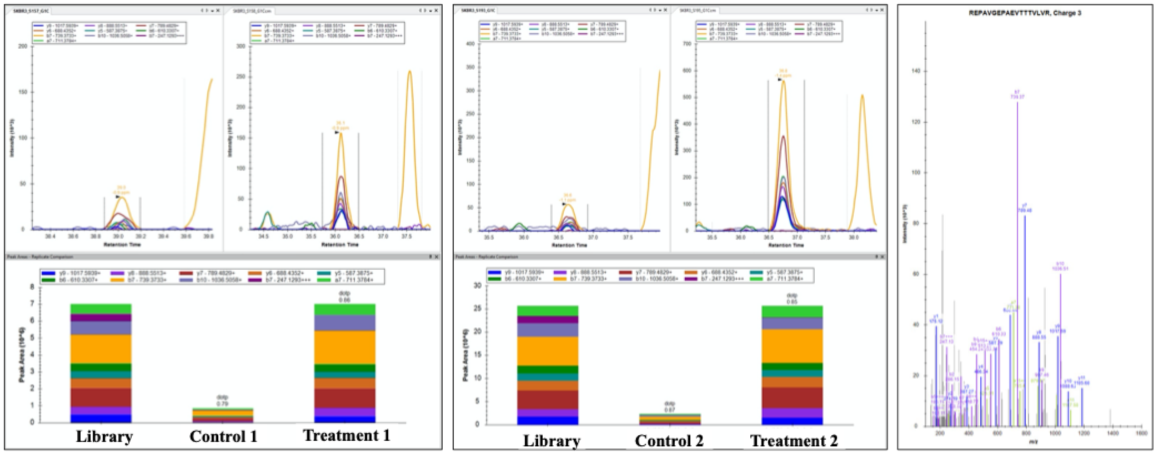


**CEBPB (CCAAT/enhancer-binding protein beta)**
**APPTACYAGAAPAPSQVK, Charge +2, m/z 850.42607**

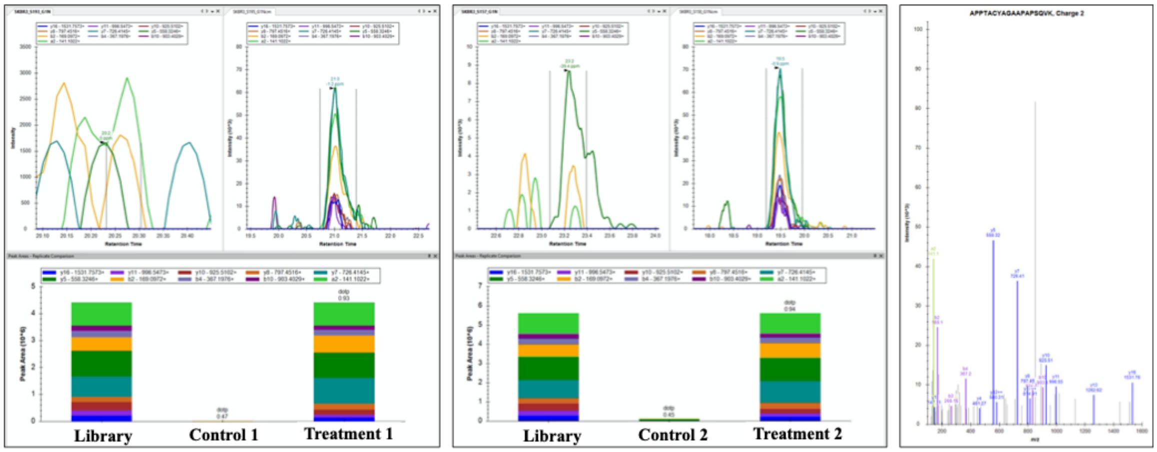

**S100A7 (Protein S100-A7)**
**IDFSEFLSLLGDIATDYHK, Charge +2, m/z 871.91786**

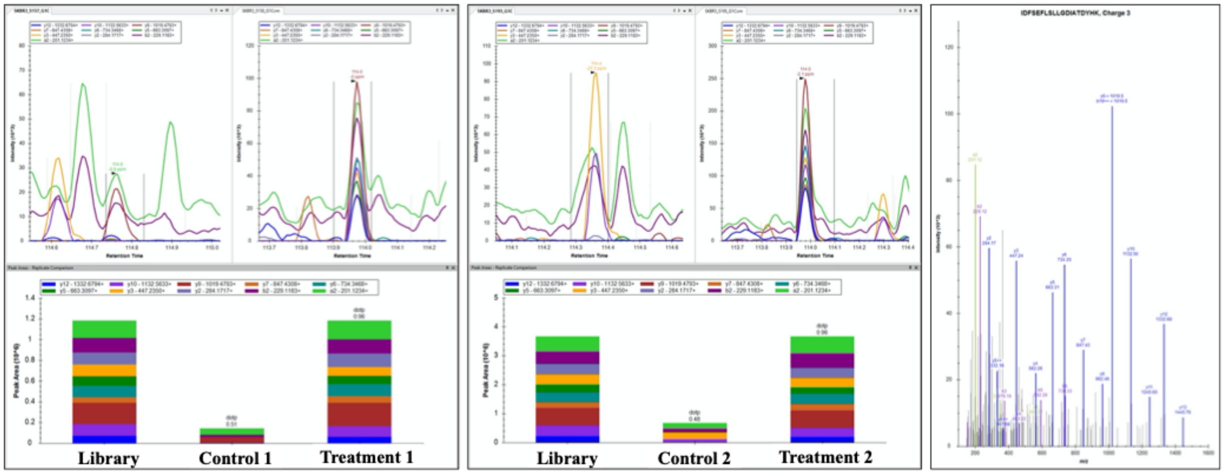


**S100A9 (Protein S100-A9)**
**VIEHIMEDLDTNADK, Charge +2, m/z 871.91786**

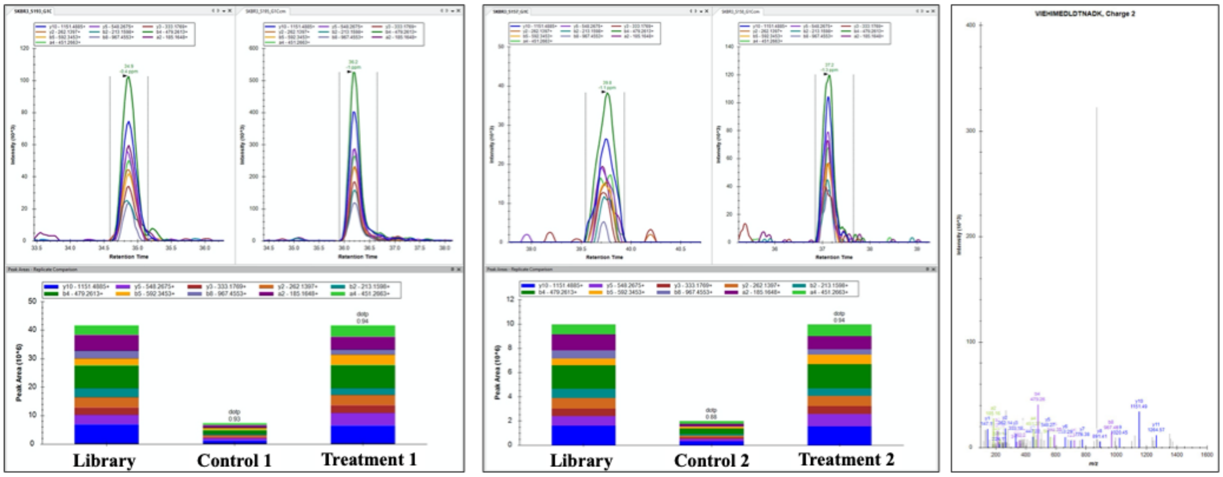


**FN1 (Fibronectin)**
**SSPVVIDASTAIDAPSNLR, Charge +2, m/z 957.00224**

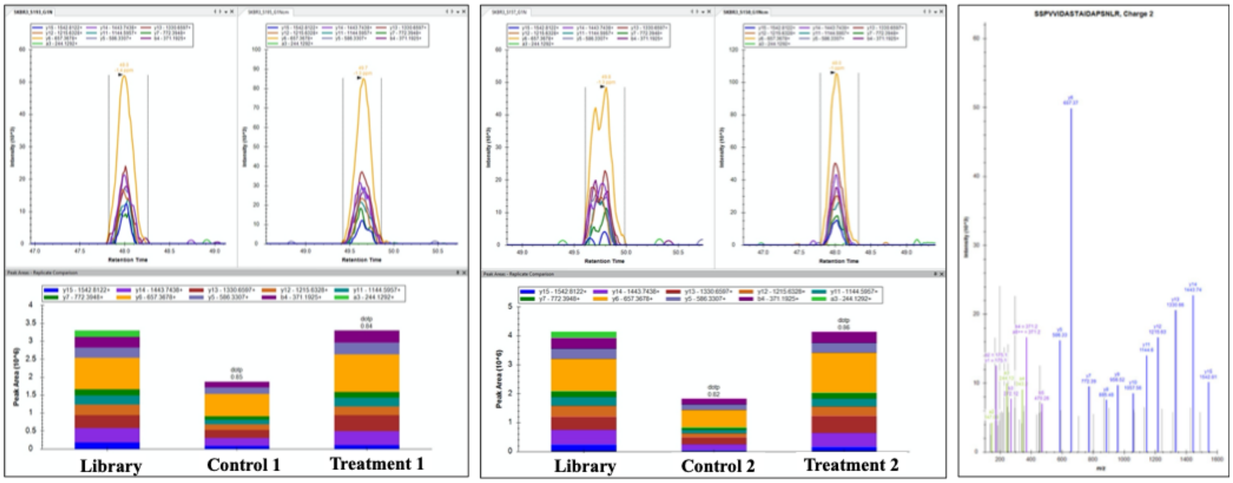

### PRM-MS validation for selected proteins with decreased counts in the CM-treated SKBR3 cells

#### CYB5A (Cytochrome b5 )
#### TFIIGELHPDDRPK, Charge +2, m/z 819.43541

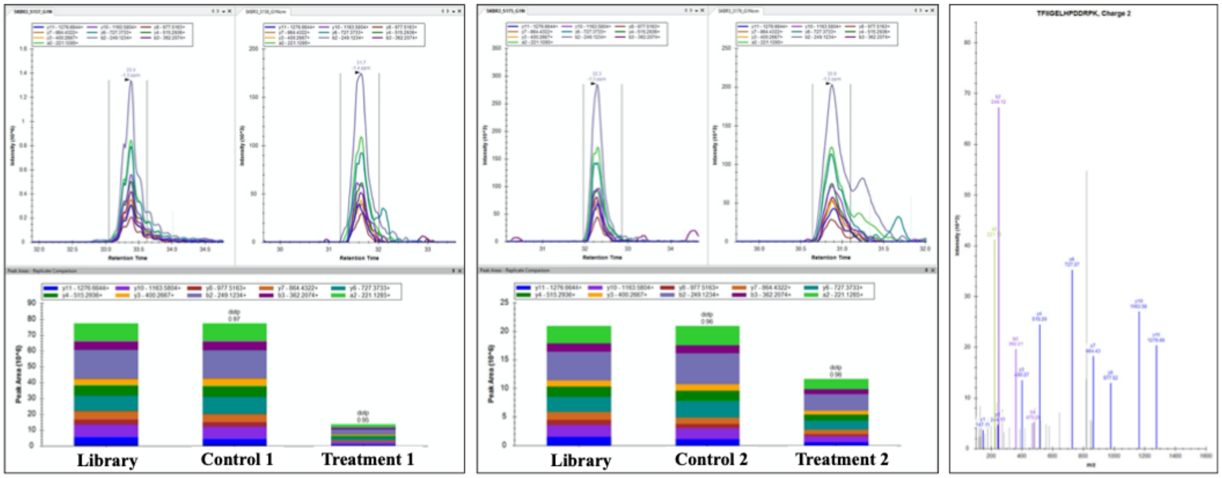


#### RPL8 (60S ribosomal protein L8)
#### ASGNYATVISHNPETK, Charge +2, m/z 844.91501

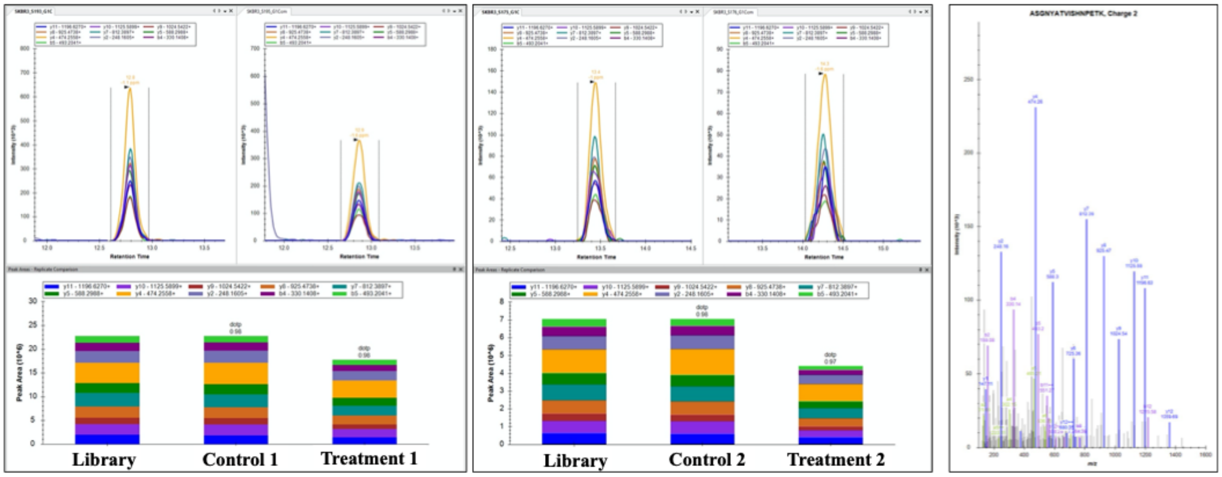


#### PRDX1 (Peroxiredoxin 1)
#### KQGGLGPMNIPLVSDPK, Charge +2, m/z 875.97854

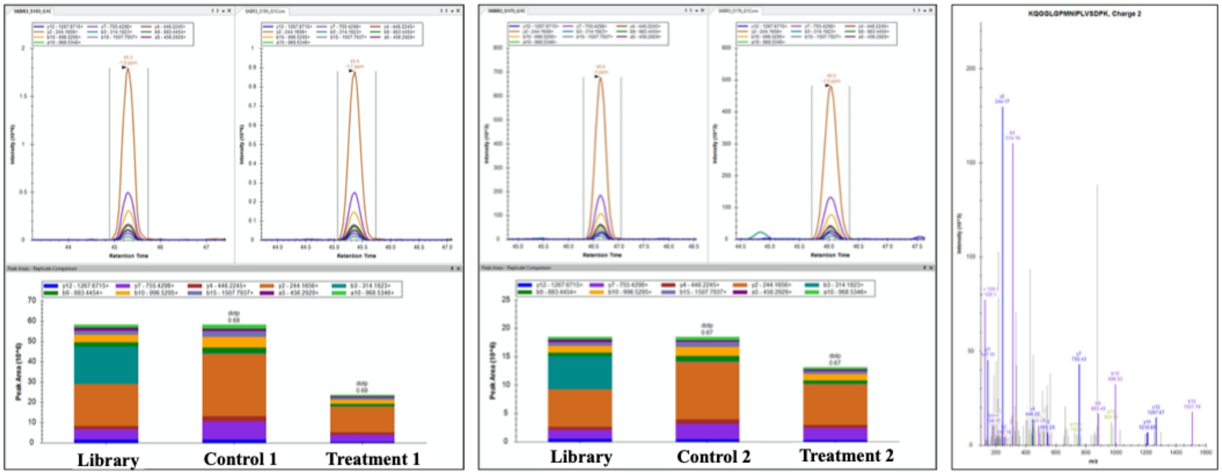

**LAMP1 (Lysosome-associated membrane glycoprotein 1)**
**FFLQGIQLNTILPDAR, Charge +2, m/z 923.51407**

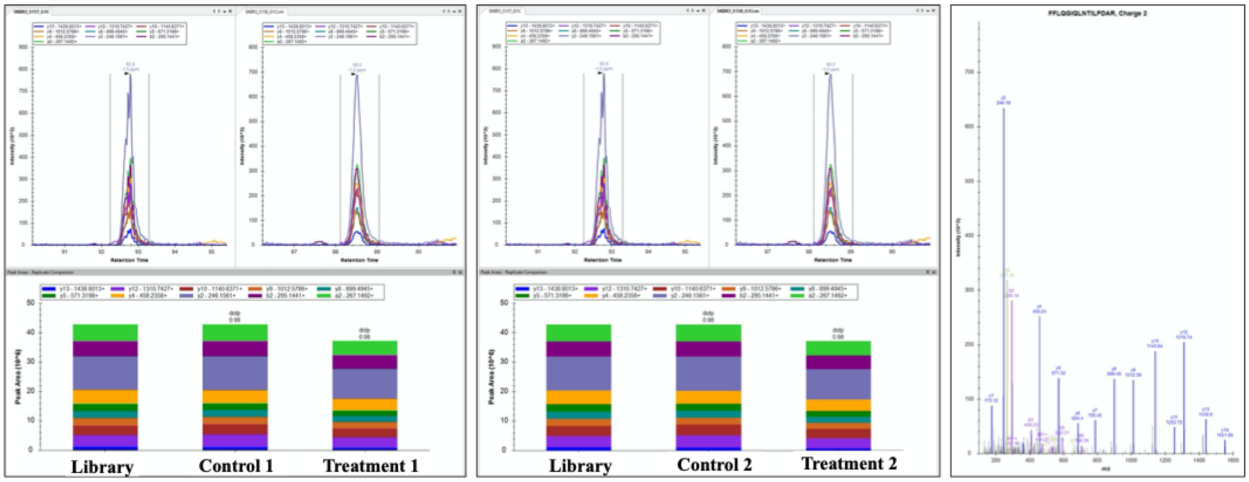


**LAT1 (Large neutral amino acids transporter small subunit 1)**
**GDVSNLDPNFSFEGTK, Charge +2, m/z 863.89885**

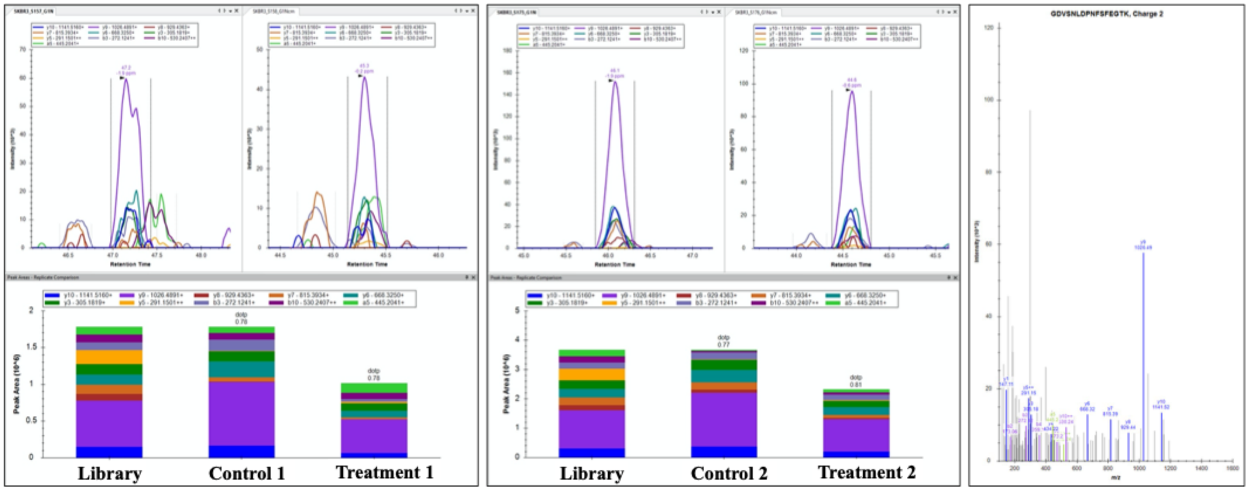


.
